# PROX1 Controls Cellular Patterning, Innervation and Differentiation in the Mouse Organ of Corti

**DOI:** 10.1101/2025.04.15.649013

**Authors:** Shubham Kale, Dimitri Traenkner, Ellison J. Goodrich, Satish R. Ghimire, Thomas M. Coate, Michael R. Deans

## Abstract

The organ of Corti is divided into functional compartments responsible for hearing or cochlear amplification. A medial compartment containing inner hair cells innervated by Type I spiral ganglion neurons and a lateral compartment containing outer hair cells innervated by Type II spiral ganglion neurons. Supporting cells also differ, with lateral compartment pillar cells and Deiters’ cells developing specialized cellular structures to support outer hair cell electromotility. We bred organ of Corti-restricted *Prox1* conditional knockout mice to study lateral compartment development because PROX1 is the first transcription factor expressed strictly in lateral compartment supporting cells. In the absence of *Prox1*, supporting cell numbers increased without corresponding changes in outer hair cells, and they appear incompletely differentiated based on morphological criteria. Outer hair cell number was not impacted but innervation was disrupted with many afferent neurons turning incorrectly towards the cochlear apex. RNAseq revealed no changes in gene expression that could account for the innervation phenotype. Therefore, we propose that PROX1 promotes pillar and Deiters’ cell differentiation and organization that has secondary effects on innervation but is not required for compartment specification.

## Introduction

The organ of Corti is the sensory epithelium in the cochlea that contains the mechanosensitive hair cells that facilitate the detection of sound and hearing. It contains two populations of hair cells and six populations of supporting cells that are precisely organized in position and number (Fig.1A). These cells can be divided along the radial axis of the cochlea into two functional compartments with a medial compartment containing the single row of inner hair cells (IHCs) that detect the soundwaves perceived as sound, and a lateral compartment containing the three rows of outer hair cells (OHCs) that comprise the cochlear amplifier. Supporting cell types are also segregated with outer pillar cells (OPC) located in the lateral compartment along with three rows of Deiters’ cells. These are positioned adjacent to the Hensen’s cells at the lateral boundary of the organ of Corti. The medial compartment contains the inner pillar cells (IPC) and the inner phalangeal cells and border cells that flank the IHCs. These compartments are separated by the tunnel of Corti that forms between the IPCs and OPCs, and both compartments spiral in parallel along the length of the cochlea (Huh, Jones et al. 2012, Driver and Kelley 2020). In addition, the hair cells in these compartments are innervated differently by afferent fibers of the cochlear nerve. Type I Spiral Ganglion Neurons (SGNs) form one-to-one connections with IHCs in the medial compartment, whereas Type II SGNs innervate multiple OHCs in the lateral compartment by extending a peripheral axon that makes a characteristic 90-degree turn toward the cochlear base. Axon turning is regulated in a non-cell autonomous manner by the asymmetric distribution of planar cell polarity (PCP) proteins localized in lateral compartment supporting cells (Ghimire, Ratzan et al. 2018, Ghimire and Deans 2019).

**Figure 1:**
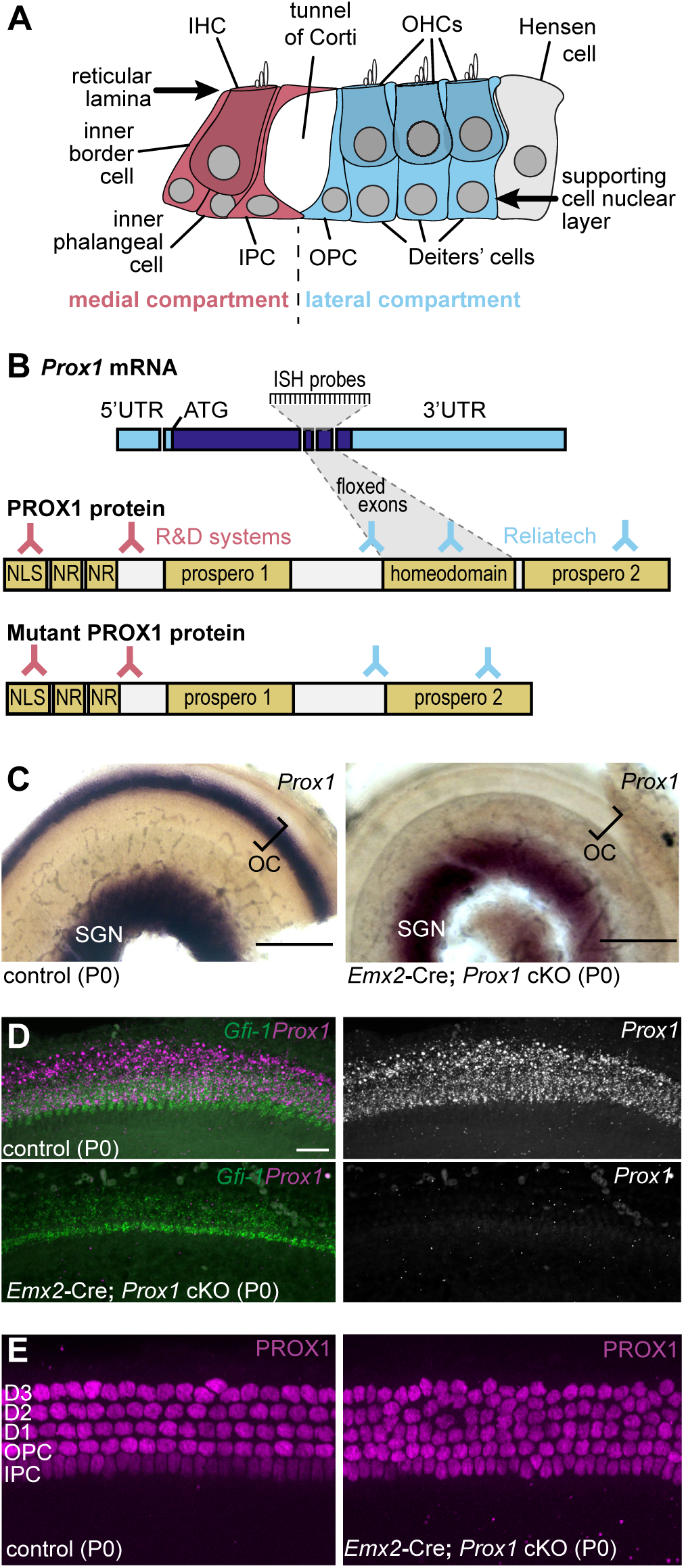
Organ of Corti restricted *Prox1* conditional knockout. (**A**) Schematic cross-section of the mouse organ of Corti identifying the medial and lateral compartments and cell types they contain. Arrows mark the position of two viewing planes used in subsequent wholemount imaging; the reticular lamina at the apical surface of hair cells and supporting cells, and a deeper plane where the supporting cell nuclei are located. (**B**) Schematic of *Prox1* mRNA depicting targeted exons, wild type and mutant PROX1 protein with described functional domains (Baxter, Cheung et al. 2011), targets of *Prox1* ISH probes and location of antigens recognized by PROX1 antibodies. Cre-mediated deletion of the targeted exons removes the homeodomain but does not prevent translation of mutant PROX1 protein or impact conserved prospero domains, nuclear receptor boxes (NR) or nuclear localization signal (NLS). (**C**) Histochemical ISH for *Prox1* mRNA at P0 demonstrates restricted gene deletion from the organ of Corti in *Emx2*-Cre; *Prox1* cKO cochleae (n=3 controls, n=4 cKOs). (**D**) HCR fISH for *Prox1* mRNA confirms exon deletion from organ of Corti in *Emx2*-Cre; *Prox1* cKOs in the vicinity of *Gfi1-*expressing hair cells (n=3 controls, n=4 cKOs). (**E**) Immunofluorescent labeling of PROX1 marks Inner Pillar Cells (IPC), Outer Pillar Cells (OPC) and three rows of Deiters cells (DC1, DC2, DC3) in littermate controls and detects mutant PROX1 protein in the cKO (n=3 controls, n=6 cKOs). Scale bars are 100 µm (C) and 20µm (D,E).

It has been proposed that the lateral compartment evolved as a mammalian specialization that enables OHC electromotility and cochlear amplification that are not characteristics of non-mammalian species (Koppl and Manley 2019). The medial and lateral compartments are distinguished during early cochlear development by morphogen gradients that establish a radial pattern across the prosensory domain that will become the organ of Corti. A WNT-signaling gradient from medial to lateral defines the medial region whereas a BMP signaling gradient from lateral to medial is required to pattern the lateral compartment (Munnamalai and Fekete 2016, Jansson, Ebeid et al. 2019). Further studies have shown that Fgf20/Fgfr1 signaling is required for outer compartment hair cell and supporting cell differentiation (Huh, Jones et al. 2012). While these morphogenetic cues determine the relative position of these compartments, separate factors must act downstream to promote compartment and cell type-specific differentiation. For example in hair cells, TBX2 is required for IHC development and fate specification (Garcia-Anoveros, Clancy et al. 2022, Kaiser, Ludtke et al. 2022) while INSM1 plays a comparable role in OHC development (Wiwatpanit, Lorenzen et al. 2018). In contrast, the transcription factors and regulatory networks that control supporting cell fate and differentiation in medial and lateral compartments are less characterized.

One candidate is the transcription factor PROX1 (Prospero homeobox gene 1) which is a member of the homeobox transcription factor family and is expressed by the IPCs, OPCs, and Deiters’ cells. Outside the cochlea, PROX1 functions to promote lymphatic endothelial cell fate, pancreas morphogenesis, and progenitor proliferation and horizontal cell specification in the retina (Hong, Harvey et al. 2002, Wigle, Harvey et al. 2002, Dyer, Livesey et al. 2003, Wang, Kilic et al. 2005). In *Drosophila*, the invertebrate ortholog PROSPERO regulates the transition between neural progenitor self-renewal and differentiation (Choksi, Southall et al. 2006), cell type specification (Doe, Chu-LaGraff et al. 1991), and axonal outgrowth (Vaessin, Grell et al. 1991). In the organ of Corti, PROX1 is expressed shortly after the sensory domain is specified by cells that will become supporting cells in the lateral compartment. This includes the IPCs which appear to have lateral identity during early development but are anatomically compartmentalized to the medial side of the tunnel of Corti after its formation. PROX1 is also expressed in spiral ganglion neurons and contributes to Type II SGNs innervation of OHCs in the lateral compartment (Fritzsch, Dillard et al. 2010). We evaluated PROX1 during organ of Corti development by generating *Prox1* conditional knockouts (cKO) in which PROX1 function is blocked in just in the organ of Corti using *Emx2*-Cre. Marker expression and transcriptional profiling show that lateral compartment cell identities remain specified in the cKO, but their differentiation is impacted with increases in their number, disrupted cellular patterning, and changes in morphology. *Prox1* is also required in the lateral compartment to guide innervation, though this appears to be secondary to the changes in supporting cell organization, and not a direct function of PROX1 in axon guidance.

## Results

### Selective Prox1 gene deletion from the mouse organ of Corti

Expression of *Prox1* begins in the basal turn of the cochlea at embryonic day 14.5 (E14.5) and progresses towards the apical turn as the cochlea matures. Initially it is expressed in hair cells, supporting cells and the spiral ganglia, but is down-regulated in hair cells and up-regulated in IPCs, OPCs and Deiters’ cells at E16.5 (Bermingham-McDonogh, Oesterle et al. 2006, Fritzsch, Dillard et al. 2010). Expression is maintained in the spiral ganglion beyond postnatal day 25 (P25) (Shrestha, Chia et al. 2018), and PROX1 has been proposed to direct turning of the Type II SGN peripheral axon towards the base of the cochlear spiral (Fritzsch, Dillard et al. 2010). To determine the contribution of *Prox1* to organ of Corti development and innervation, a conditional knockout was generated by crossing *Emx2*-Cre knock-in mice (Kimura, Suda et al. 2005) with a *Prox1* floxed line in which LoxP sites flank two exons encoding the PROX1 homeobox domain (Fig.1B) (Harvey, Srinivasan et al. 2005). These correspond to exons 3 and 4 based upon a current GenBank entry for *Prox1* RNA (NM_008937.3) In the resulting *Emx2* ^Cre/wt^; *Prox1* ^LoxP/LoxP^ mice (henceforth *Emx2*-Cre; *Prox1* cKO) Cre-mediated recombination is restricted to the cochlear duct, occurring at E12.5 (Ono, Kita et al. 2014, Goodrich and Deans 2024) and before the onset of *Prox1* expression. Effective gene targeting was assessed by *in-situ* hybridization (ISH) using conventional histochemical techniques (Fig.1C) and an HCR probe for fluorescent ISH using the Hybridization Chain Reaction (HCR) technique (Fig.1D) (Choi, Schwarzkopf et al. 2018). Both probes are specific for the targeted exons (Fig.1B) and demonstrate deletion of those exons from the cochlear duct but not the SGN. The loss of these exons from *Prox1* causes an in-frame deletion and expression of a mutant PROX1 protein that remains stable but is missing DNA binding domains. Notably, this leaves the antigens recognized by two different PROX1 antibodies intact (Fig.1B), and mutant PROX1 can be visualized in the cKO by immunofluorescent labeling (Fig.1E). As previously reported, PROX1 is expressed by IPCs, OPCs and the three rows of Deiters’ cells at postnatal day 0 (P0), where it localizes to the nucleus (Bermingham-McDonogh, Oesterle et al. 2006, Fritzsch, Dillard et al. 2010). Mutant PROX1 has the same distribution in *Emx2*-Cre; *Prox1* cKOs and as a result, its expression can be used as a marker to follow the development of lateral compartment supporting cells in the absence of functional PROX1 protein.

### Deletion of Prox1 results in increased numbers of Supporting Cells

A prominent feature of the *Emx2*-Cre; *Prox1* cKO phenotype are cellular patterning defects that disrupt the organized rows of supporting cell nuclei. This is readily observed when wholemount *Emx2*-Cre; *Prox1* cKO cochleae are immunolabeled for mutant PROX1 protein (Fig.1E, 2A). This disorganization does not occur in *Emx2*-Cre; *Prox1* ^LoxP/wt^ cochleae indicating that mutant PROX1 does not have a dominant negative effect. Therefore, to minimize variables, only Cre-positive tissues were used for analysis with *Emx2* ^Cre/wt^; *Prox1* ^LoxP/wt^ serving as littermate controls. Quantification reveals that the disorganization is due to the presence of extra supporting cells expressing mutant PROX1 (Fig.2B). This phenotype was not evident at E17.5, but appeared at P0 and then remained constant for the ensuing seven days of postnatal development (Fig.2A,B). When evaluated in sections at E17.5, the ratio of supporting cells to hair cells did not differ between *Emx2*-Cre; *Prox1* cKOs and littermate controls consistent with the PROX1-labeled cell quantification in wholemounts at this age (Fig.2C). Thus, between E17.5 and P0 there was on average a 24% increase in the number of PROX1 immunolabeled nuclei in *Emx2*-Cre; *Prox1* cKOs compared to littermate controls and this occurred at all positions along the length of the cochlear spiral (Fig.2B). The extra cells were not organized into a discrete row and instead were interspersed between the other OPCs and Deiters’ cells. A similar observation of disorganized nuclei was observed in *Prox1* cKOs generating using *Pax2*-Cre and evaluated by transmission electron microscopy (Fritzsch, Dillard et al. 2010). In comparison there were no changes in the number of hair cells in the *Emx2*-Cre; *Prox1* cKO per unit length (Fig.2D) and the additional supporting cells did not impact the overall length of the cochlear spiral (Fig.2E).

**Figure 2:**
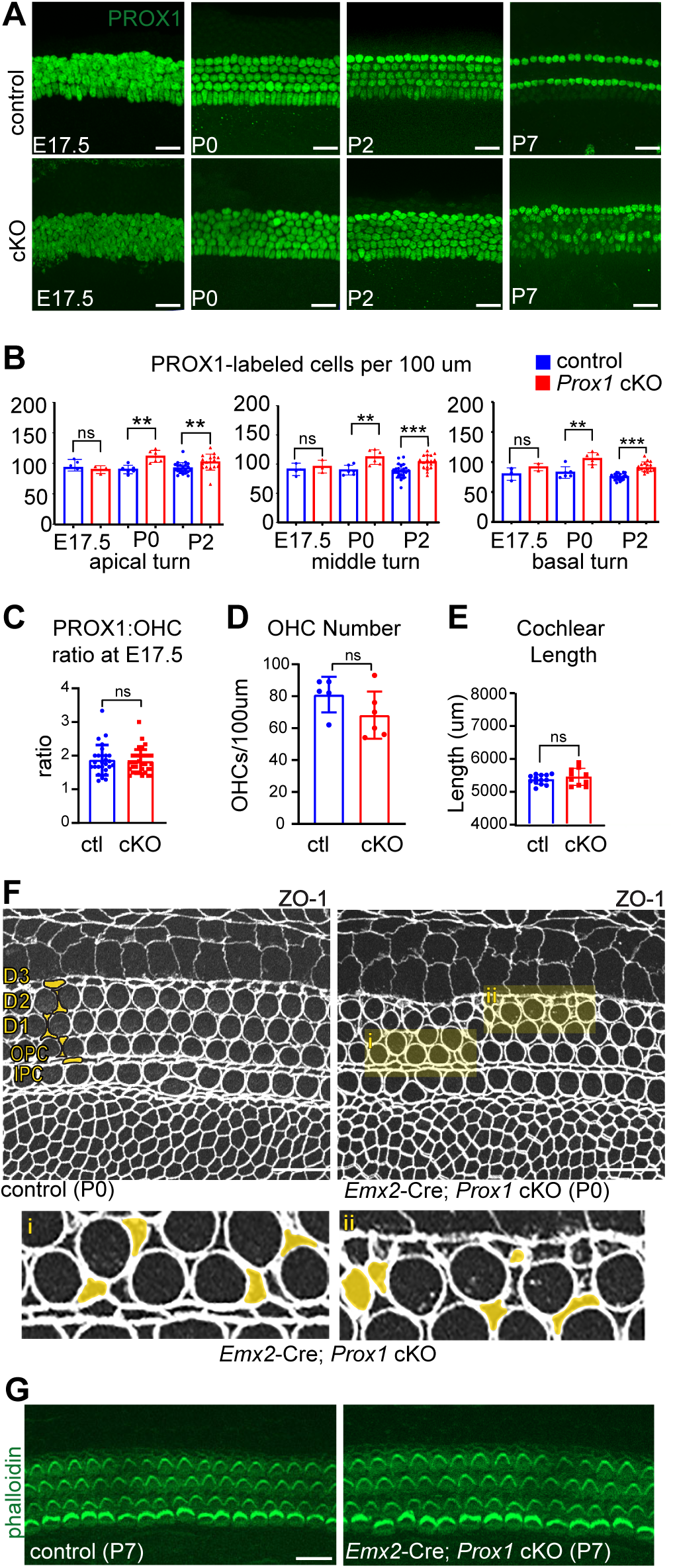
*Emx2-*Cre; *Prox1* cKOs produce extra supporting cells. (**A**) Wholemount immunofluorescent labeling of PROX1 or mutant PROX1 in *Emx2-*Cre; *Prox1* cKOs and littermate controls during neonatal and postnatal development. All images were taken from the middle turn. (**B**) The number of PROX1 immunolabeled nuclei per 100µm of cochlear length at neonatal stages and at three positions along the cochlear spiral (n=4 (E17.5), 6 (P0), 22 (P2) for controls and (n=3 (E17.5), 6 (P0), 22 (P2) for cKOs). (**C**) The ratio of supporting cells to hair cells at E17.5 does not differ between cKOs (n=4 cochleae, 29 sections) and controls (n=4 cochleae, 26 sections) when evaluated in sectioned cochleae. (**D**) The number of OHCs per 100µm of cochlear length does not differ between cKOs (n=6) and controls (n=5) at P0. (**E**) The overall length of the cochlear spiral at P2 does not change between cKOs (n=10) and controls (n=13). (**F**) ZO-1 immunolabeling of the reticular lamina shows the distribution of Deiters’ and OPC processes between OHCs. Higher magnification fields (**i, ii, iii**) from the *Emx2*-Cre; *Prox1* cKO where extra supporting cell processes are marked by colored shading. (**G**) Phalloidin stain of wholemount cochleae shows no difference in hair cell distribution, bundle morphology or polarity between cKOs (n=6) and littermate controls (n=5). Error bars show SEM, and asterisks indicate statistical significance for differences between genotypes using Student’s *t*-test (**p<0.01, ***p<0.001). Scale bars are 20µm.

Deiters’ cells are unique because their cell somas are positioned beneath the OHCs and they project a phalangeal process that extends diagonally towards the cochlear apex before contacting the reticular lamina. The apical surface of the phalangeal processes are precisely patterned with one supporting cell located between two neighboring hair cells in each row. This pattern can be visualized by immunolabeling the tight junctions of the reticular lamina with antibodies against ZO-1 (Fig.2F). Using this approach, the circular outline of hair cells are readily identified in control cochleae where they are precisely aligned in three rows with dumbbell shaped supporting cell outlines in between. While both cell types are also outlined in the *Emx2*-Cre; *Prox1* cKO, ZO-1 labeling also reveals apical projections from the additional supporting cells interspersed between OHCs. These ectopic projections occurred within all three OHC rows with a frequency and distribution similar to the extra supporting cell nuclei expressing mutant PROX1. In contrast, the distribution and planar polarization of hair cells was not impacted by the loss of *Prox1* (Fig.2G).

#### Extra supporting cells are not correlated with increased proliferation or apoptosis

Supporting cells are normally born during a period of mitosis occurring between E14 and E15.5 throughout the pro-sensory domain. The timing of these cell divisions parallels that of hair cells, and no additional mitoses of either cell type occurs after E16 (Ruben 1967). The possibility that extra supporting cells appearing between E17.5 and P0 were due to ectopic proliferation at this stage was evaluated in *Emx2*-Cre; *Prox1* cKOs by EdU birth dating experiments. EdU was administered to timed pregnant mice twice daily on days E16.5, E17.5, and E18.5 of gestation and cochleae were collected and analyzed by immunofluorescent labeling for PROX1 and EdU. While proliferating cells outside of the organ of Corti were labeled, EdU incorporation did not overlap with mutant PROX1 or PROX1 expressing cells in *Emx2*-Cre; *Prox1* cKOs or littermate controls (Fig.3A). It was also not the case that the normal period of proliferation was expanded because labeling for the proliferating cell marker Ki-67 did not mark Prox1-expressing cells at E15.5 (Fig.3B). Together these results were not consistent with additional rounds of cell division occurring in the *Emx2*-Cre; *Prox1* cKO. Alternatively, additional supporting cells may be present in the cKO due to decreases in supporting cell apoptosis. Although apoptosis is not a significant feature of cochlear development, this possibility was tested by analyzing *Bax* knockout mice where apoptosis does not occur. Immunolabeling for mutant PROX1 revealed no additional supporting cells in *Bax* knockouts, and the ratio of supporting cells to OHCs and the organized pattern of PROX1-labeled nuclei unchanged between knockouts and controls (Fig.3C).

**Figure 3:**
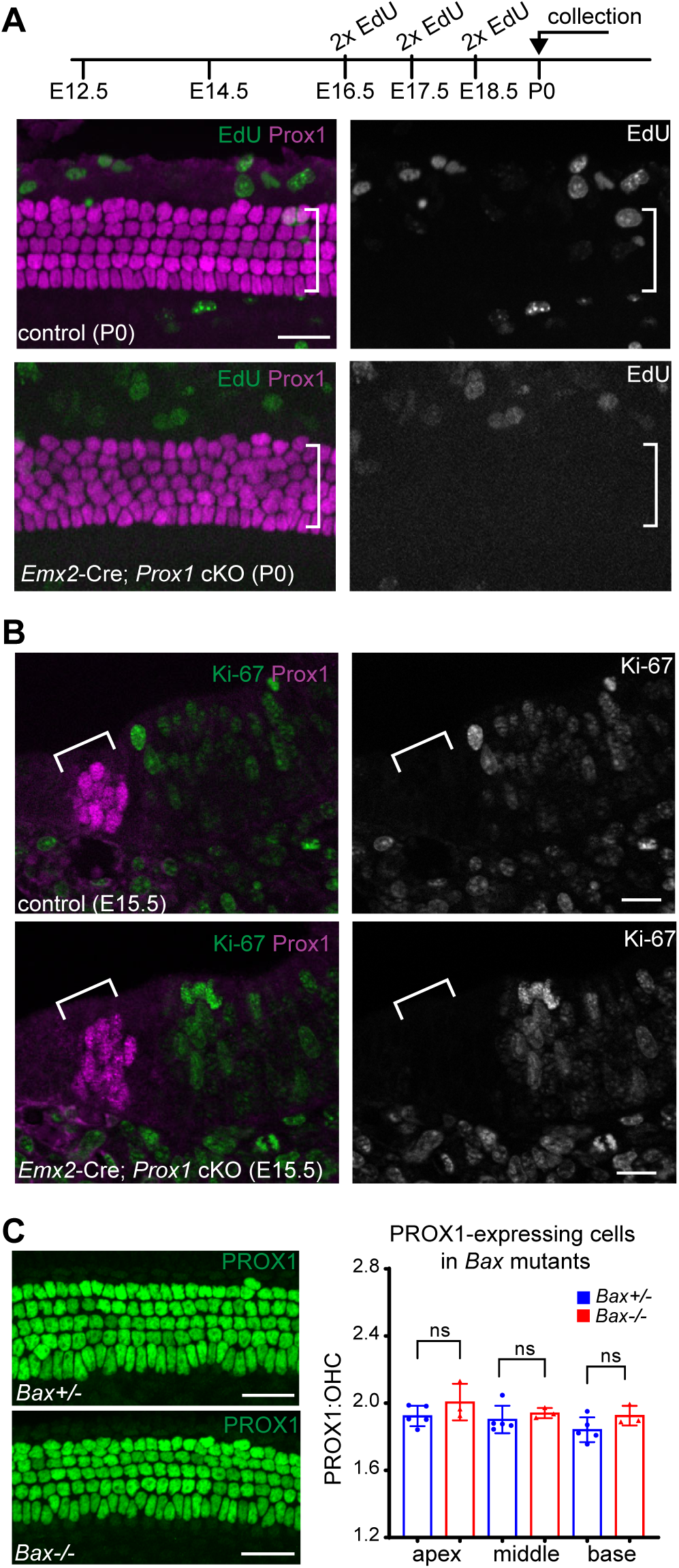
Extra supporting cells are not due to increased mitosis or the absence of apoptosis. (**A**) Experimental timecourse of EdU delivery to timed pregnant dams and wholemount immunolabeling of P0 cochleae for EdU-labeled and PROX1 or mutant PROX1 expressing cells in *Emx2-*Cre; *Prox1* cKOs (n=4) and littermate controls (n=3). (**B**) Immunolabeling for the cell division marker Ki-67 shows no overlap with PROX1 or mutant PROX1 expressing cells in sectioned cochleae at E15.5 from *Emx2-*Cre; *Prox1* cKOs (n=4) or littermate controls (n=5). (**C**) Wholemount immunolabeling for PROX1 in the *Bax* knockout (*Bax* ^-/-^, n=3) or heterozygous control (*Bax ^+/-^,* n=5) cochleae and ratios of PROX1-labeled cells to OHCs at three positions along the length of the cochlear spiral. Error bars show standard deviation, and asterisks indicate statistical significance for differences between genotypes using Student’s *t*-test (**p<0.01, ***p<0.001). Brackets in A&B mark the boundaries of the PROX1-expressing domains. Scale bars are 20µm (A,C) and 10µm (B).

#### Disrupted Type II SGN innervation patterns in the *Emx2*-Cre; *Prox1* cKOs

*Prox1* was previously shown to be necessary for peripheral axons of the Type II SGNs to turn towards the cochlear base (Fritzsch, Dillard et al. 2010) and PROX1 is expressed by both organ of Corti supporting cells and SGNs. However, when evaluated by immunofluorescent labeling, PROX1 levels decrease in the Type II SGNs during postnatal development (Nishimura, Noda et al. 2017), and transcriptomic evaluation of SGNs showed greater and sustained expression of *Prox1* in Type I SGNs compared to Type II SGNs (Petitpre, Wu et al. 2018). Together these expression assays raise the possibility that *Prox1* may not act autonomously within Type II SGNs to regulate axon guidance. Alternatively, the disorganized pattern of supporting cells in the *Emx2*-Cre; *Prox1* cKO suggest changes in the environment traversed by the peripheral axons may be the cause of growth cone turning errors. This possibility was evaluated by comparing the Type II SGN innervation in the *Emx2*-Cre, *Prox1* cKO with a *Bhlhe22*-Cre; *Prox1* cKO where Cre activity is restricted to the developing SGN (Appler, Lu et al. 2013). The efficiency of *Bhlhe22*-Cre mediated gene deletion was demonstrated by conventional histochemical ISHs against the targeted *Prox1* exons (Fig.4A). Despite the efficiency of *Prox1* gene deletion, axon pathfinding defects were not evident in *Bhlhe22*-Cre; *Prox1* cKOs but were readily apparent in *Emx2*-Cre; *Prox1* cKOs. (Fig.4B,C). The incorrectly turning axons in cochlear duct restricted cKO reveal a non-cell autonomous function for PROX1 in the lateral compartment supporting cells to direct axon guidance and OHC innervation.

**Figure 4:**
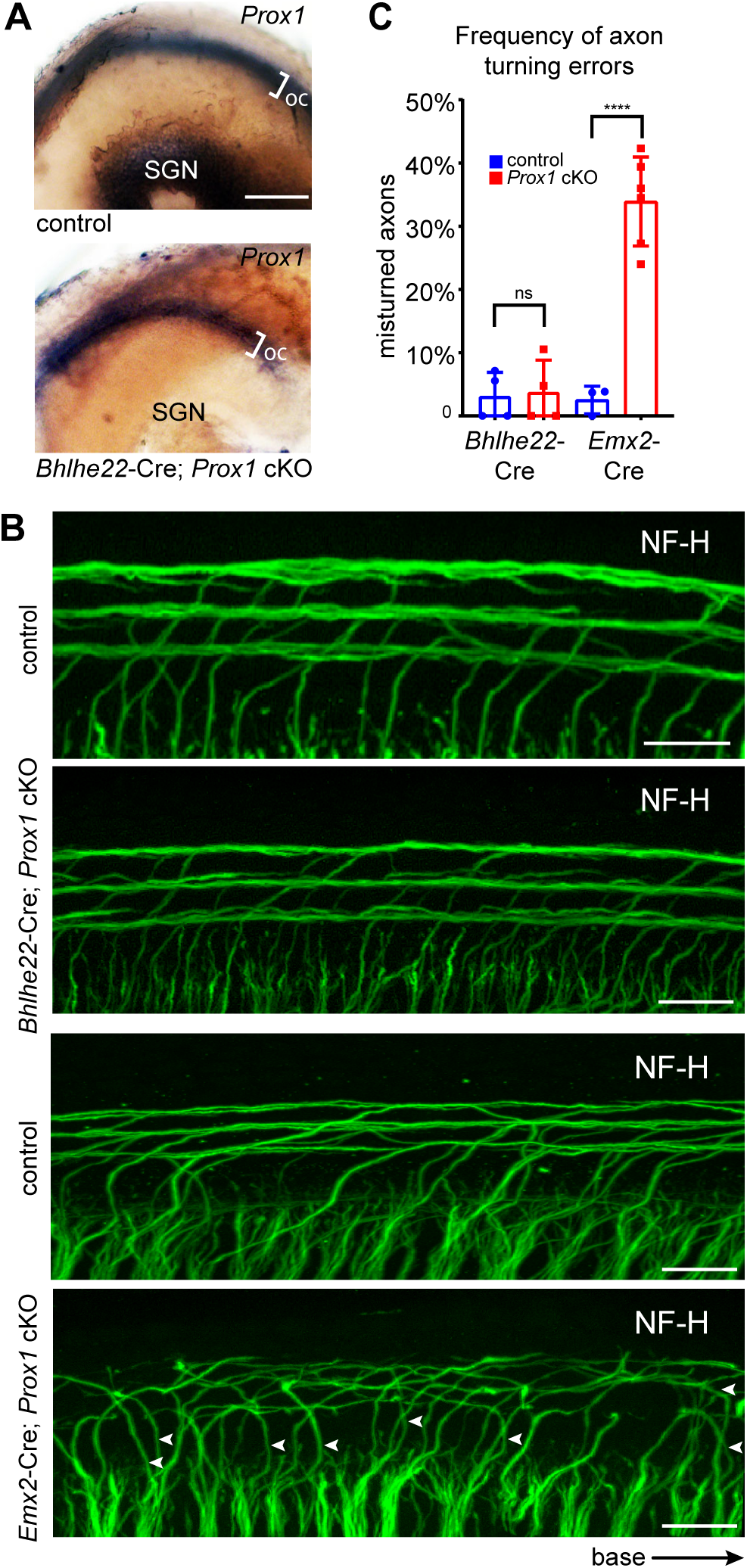
PROX1 directs Type II SGN axon turning non-cell autonomously. (**A**) Conventional histochemical ISH of *Bhlhe22-*Cre; *Prox1* cKO (n=3) and littermate control (n=3) cochleae using probes against the targeted *Prox1* exons to assess expression in the organ of Corti (OC) and spiral ganglion neurons (SGN). (**B**) Peripheral axons of Type II SGNs immunolabeled for Neurofilament (NF-H) in *Emx2-*Cre; *Prox1* cKOs (n=6) or littermate controls (n=3), and *Bhlhe22-*Cre; *Prox1* cKOs (n=4) and littermate controls (n=4). Arrowheads indicate incorrectly turning axons. (**C**) Quantification of axonal turning errors for Type II SGNs located in the basal turn of the cochlea. Error bars show standard deviation, and asterisks indicate statistical significance for differences between genotypes using Student’s *t*-test (****p<0.0001). Scale bars are 100µm (A) and 20µm (B).

#### RNAseq quantification of axon guidance gene expression *Prox1* cKOs

As a transcription factor, PROX1 may function non-cell autonomously by regulating the expression of axon guidance cues in the developing organ of Corti at the time when peripheral axons are turning towards the cochlear base. PCP signaling has also been shown to direct peripheral axon turning, and *Vangl2* and *Fzd3/Fzd6* conditional knockouts generated using *Emx2*-Cre have innervation phenotypes that resemble the *Emx2-*Cre; *Prox1* cKO (Ghimire, Ratzan et al. 2018, Ghimire and Deans 2019). More recently, the cytoskeletal regulator Rac1 and the cell adhesion molecule NECTIN3 have also been shown to regulate axon turning in a non-cell autonomous manner (Clancy, Xie et al. 2025). To determine whether PROX1 regulates PCP or axon guidance gene expression, bulk RNA sequencing was used to measure changes in gene expression between *Emx2*-Cre; *Prox1* cKOs and littermate controls at P0. Cochleae were dissected to separate the organ of Corti from the SGN, and organ of Cortis from multiple pups were pooled for RNA extraction, library prep and sequencing. Differentially expressed genes (DEG) were identified, and 50 genes were down-regulated and 36 genes were up-regulated in the cKO (Fig.5A and Tables 1&2). When compared to available single-cell RNAseq datasets from postnatal mouse cochleae (Kolla, Kelly et al. 2020) it was apparent that the expression patterns of many DEGs were not restricted to *Prox1*-expressing supporting cells but also included hair cells which transiently express PROX1 during their development (Bermingham-McDonogh, Oesterle et al. 2006), and other cell types not expressing *Prox1* (Fig.5B). Due to the mechanical methods used to isolate the organ of Corti these may include mesenchymal and other non-sensory cell types. However, when the relative contribution of DEG transcripts to the overall transcriptional profiles of these groups was evaluated, it became apparent that the down-regulated genes were genes normally found at higher levels in *Prox1*-expressing supporting cells than the other groups (Fig.5B). In contrast, most genes that were up-regulated were ones normally expressed in all three groups. An exception was *Prox1,* which showed a 0.32 log2fold overall increase by DESeq2 (Supp. Table2), with a 25% increase in sequencing reads that aligned with non-targeted exons despite the expected reduction in reads aligning to targeted exons (Fig.5C). This is likely due to the increase in *Prox1*-expressing cells but may also reflect increased transcription from the *Prox1* promoter in cKOs. Another upregulated gene was *Atb8b4* which only appeared in a small number of cochlear cells in the scRNAseq dataset used to create these groups and therefore was not identified or included in this analysis.

**Figure 5:**
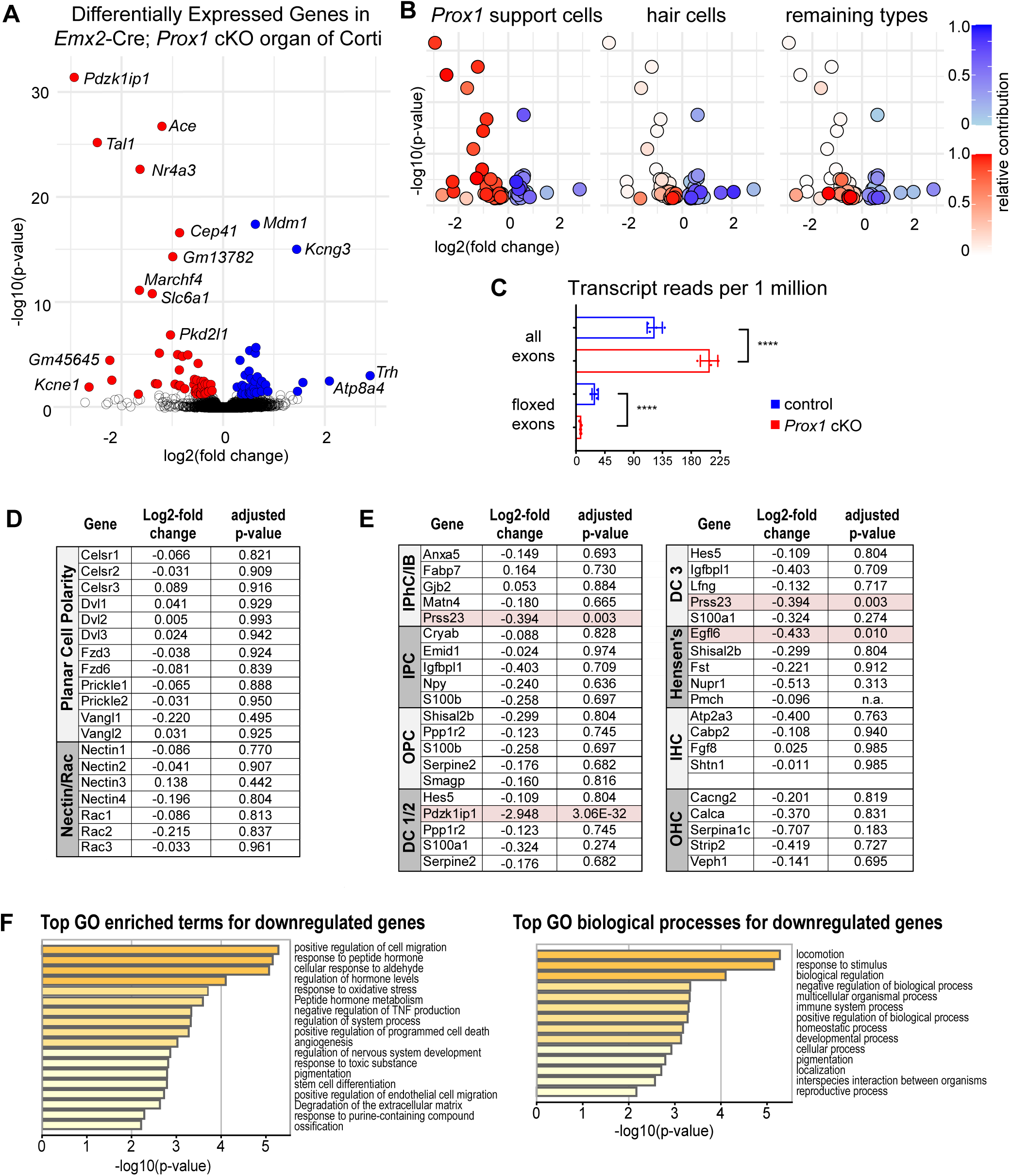
RNA-seq analysis of *Emx2-*Cre; *Prox1* cKOs. (**A**) Differentially expressed genes (DEG) between *Emx2*-Cre; *Prox1* cKO and littermate control organ of Corti at P0 identified using DESeq2 with genes significantly downregulated in the cKO in red and those upregulated in blue. (**B**) Volcano plots restricted to the DEGs found in different groups of cochlear cell types with color density indicating the predicted contribution of that gene to the group’s overall transcriptome. Cell type groups and levels of gene expression are derived from public scRNAseq datasets generated from wild type tissue (Kolla, Kelly et al. 2020). (**C**) Average values of normalized *Prox1* transcript alignments against all *Prox1* exons compared to alignments against the two exons targeted in *Emx2*-Cre; *Prox1* cKOs (n=4) and littermate controls (n=4). Error bars show standard deviation, and asterisks indicate statistical significance using Student’s *t*-test (****p<0.0001). (**D**) Changes in gene expression determined by DSeq2 for PCP, Nectin and Rac genes. (**E**) Changes in expression for genes signifying medial compartment and lateral compartment cell types. Red shading marks DEGs with statistically significant changes between genotypes (see also Supp. Table1&3). (**F**) Metascape Gene Ontology (GO) analysis of downregulated genes in the organ of Corti of *Emx2-*Cre; *Prox1* cKOs.

**Table 1:** Genes with decreased transcription in the *Emx2*-Cre; *Prox1* cKO organ of Corti. List of all 50 genes that were down-regulated and had decreased transcription in the *Emx2*-Cre; *Prox1* cKO compared to littermate controls based. Changes were considered significant for adjusted p-values less than 0.05. Genes are order based upon log2-fold change in expression.

| Ensemble ID | Gene Name | Description | log2 Fold Change | p-value | adjusted p-value |
| --- | --- | --- | --- | --- | --- |
| ENSMUSG00000028716 | *Pdzk1ip1* | PDZK1 interacting protein 1 | -2.94777 | 1.6033E-36 | 3.06E-32 |
| ENSMUSG00000032293 | *Kcne1* | potassium voltage-gated channel, Isk-related subfamily, member 1 | -2.65348 | 2.6282E-05 | 9.66E-03 |
| ENSMUSG00000028717 | *Tal1* | T cell acute lymphocytic leukemia 1 | -2.49049 | 7.8705E-30 | 5.01E-26 |
| ENSMUSG000000109849 | *Gm45645* | Predicted gene 45645 | -2.24344 | 2.5957E-08 | 2.61E-05 |
| ENSMUSG00000033470 | *Cysltr2* | cysteinyl leukotriene receptor 2 | -2.20633 | 3.3263E-06 | 2.14E-03 |
| ENSMUSG00000020660 | *Pomc* | pro-opiomelanocortin-alpha | -1.68567 | 2.0018E-04 | 4.55E-02 |
| ENSMUSG00000039372 | *Marchf4* | Membrane associated ring-CH-type finger 4 | -1.66197 | 2.7357E-15 | 5.81E-12 |
| ENSMUSG00000028341 | *Nr4a3* | nuclear receptor subfamily 4, group A, member 3 | -1.65088 | 3.7289E-27 | 1.78E-23 |
| ENSMUSG00000030310 | *Slc6a1* | solute carrier family 6 (neurotransmitter transporter, GABA), member 1 | -1.40796 | 6.1735E-15 | 1.18E-11 |
| ENSMUSG00000040148 | *Hmx3* | H6 homeobox 3 | -1.35045 | 8.9031E-06 | 4.60E-03 |
| ENSMUSG00000007480 | *Mc5r* | melanocortin 5 receptor | -1.31087 | 9.3339E-06 | 4.69E-03 |
| ENSMUSG00000046318 | *Ccbe1* | collagen and calcium binding EGF domains 1 | -1.27166 | 4.4944E-09 | 5.73E-06 |
| ENSMUSG00000020681 | *Ace* | angiotensin I converting enzyme (peptidyl-dipeptidase A) 1 | -1.22265 | 1.4621E-31 | 1.40E-27 |
| ENSMUSG00000037578 | *Pkd2l1* | polycystic kidney disease 2-like 1 | -1.05205 | 6.1098E-11 | 1.06E-07 |
| ENSMUSG000000082516 | *Gm13782* | predicted gene 13782 | -1.00862 | 1.5617E-18 | 3.73E-15 |
| ENSMUSG00000020688 | *Pmel* | premelanosome protein | -0.96224 | 1.0361E-05 | 5.08E-03 |
| ENSMUSG00000015863 | *Frzb* | frizzled-related protein | -0.90980 | 6.6631E-09 | 7.96E-06 |
| ENSMUSG00000078592 | *Mansc4* | MANSC domain containing 4 | -0.88373 | 1.9062E-05 | 7.43E-03 |
| ENSMUSG00000005608 | *Bean1* | brain expressed, associated with Nedd4, 1 | -0.87932 | 2.5751E-07 | 2.24E-04 |
| ENSMUSG00000038344 | *Cep41* | centrosomal protein 41 | -0.87237 | 6.2299E-21 | 1.98E-17 |
| ENSMUSG00000029371 | *Aldh1a7* | aldehyde dehydrogenase family 1, subfamily A7 | -0.86394 | 1.5111E-05 | 6.28E-03 |
| ENSMUSG00000029505 | *Aldh1a1* | aldehyde dehydrogenase family 1, subfamily A1 | -0.81764 | 1.0798E-08 | 1.15E-05 |
| ENSMUSG00000037663 | *Pkp1* | plakophilin 1 | -0.77263 | 4.1794E-05 | 1.43E-02 |
| ENSMUSG00000090084 | *4930523C07Rik* | RIKEN cDNA 4930523C07 gene | -0.70896 | 7.4219E-09 | 8.34E-06 |
| ENSMUSG00000021356 | *Dgat2* | diacylglycerol O-acyltransferase 2 | -0.58973 | 1.1694E-05 | 5.32E-03 |
| ENSMUSG00000024621 | *Anxa1* | annexin A1 | -0.57808 | 2.6155E-06 | 1.79E-03 |
| ENSMUSG00000032418 | *Mgp* | matrix Gla protein | -0.56577 | 6.5629E-05 | 2.13E-02 |
| ENSMUSG00000038345 | *Exph5* | exophilin 5 | -0.56456 | 1.2939E-04 | 3.39E-02 |
| ENSMUSG00000026365 | *Cfh* | complement component factor h | -0.53697 | 1.6080E-04 | 3.94E-02 |
| ENSMUSG00000022196 | *Lbp* | lipopolysaccharide binding protein | -0.53525 | 2.0402E-04 | 4.59E-02 |
| ENSMUSG00000023056 | *Car3* | carbonic anhydrase 3 | -0.53212 | 4.7397E-05 | 1.59E-02 |
| ENSMUSG00000019800 | *Cemip* | cell migration inducing protein, hyaluronan binding | -0.52952 | 3.3521E-06 | 2.14E-03 |
| ENSMUSG00000038346 | *Arhgap27* | Rho GTPase activating protein 27 | -0.50869 | 5.8571E-08 | 5.33E-05 |
| ENSMUSG00000023057 | *Pla2g7* | phospholipase A2, group VII (platelet-activating factor acetylhydrolase, plasma) | -0.49755 | 1.0111E-04 | 2.84E-02 |
| ENSMUSG00000038347 | *Cntnap4* | contactin associated protein-like 4 | -0.46224 | 8.5406E-05 | 2.55E-02 |
| ENSMUSG00000038348 | *Lmcd1* | LIM and cysteine-rich domains 1 | -0.45507 | 1.7418E-04 | 4.13E-02 |
| ENSMUSG00000001506 | *Col1a1* | collagen, type I, alpha 1 | -0.45151 | 1.7353E-04 | 4.13E-02 |
| ENSMUSG00000006775 | *Nos1* | nitric oxide synthase 1, neuronal | -0.44421 | 4.5922E-06 | 2.67E-03 |
| ENSMUSG00000038349 | *Egfl6* | EGF-like-domain, multiple 6 | -0.43349 | 2.9044E-05 | 1.05E-02 |
| ENSMUSG00000038350 | *Thbs2* | thrombospondin 2 | -0.42056 | 1.6048E-05 | 6.53E-03 |
| ENSMUSG00000038351 | *Prss23* | protease, serine 23 | -0.39416 | 4.7155E-06 | 2.67E-03 |
| ENSMUSG00000090085 | *2310022B05Rik* | RIKEN cDNA 2310022B05 gene | -0.38314 | 8.9959E-05 | 2.64E-02 |
| ENSMUSG00000038352 | *Nell2* | NEL-like 2 | -0.38023 | 1.7495E-04 | 4.13E-02 |
| ENSMUSG00000078669 | *Ctsk* | cathepsin K | -0.36716 | 1.1021E-05 | 5.14E-03 |
| ENSMUSG00000038353 | *Pdgfrl* | platelet-derived growth factor receptor-like | -0.35696 | 1.5409E-04 | 3.86E-02 |
| ENSMUSG00000060803 | *Gstp1* | Glutathione S-transferase, pi 1 | -0.31940 | 1.0702E-05 | 5.11E-03 |
| ENSMUSG00000038355 | *Sema5a* | sema domain, seven thrombospondin repeats (type 1 and type 1-like), transmembrane domain (TM) and short cytoplasmic domain, (semaphorin) 5A | -0.30146 | 7.8952E-05 | 2.40E-02 |
| ENSMUSG00000038356 | *Rassf2* | Ras association (RalGDS/AF-6) domain family member 2 | -0.29845 | 9.1112E-05 | 2.64E-02 |
| ENSMUSG00000005886 | *Kit* | kit oncogene | -0.25691 | 1.2364E-05 | 5.50E-03 |
| ENSMUSG00000026641 | *Nid1* | nidogen 1 | -0.23395 | 6.5216E-05 | 2.13E-02 |

**Table 2:** Genes with increased transcription in the *Emx2*-Cre; *Prox1* cKO organ of Corti. List of all 36 genes that were up-regulated and had increased transcription in the *Emx2*-Cre; *Prox1* cKO compared to littermate controls based. Changes were considered significant for adjusted p-values less than 0.05. Genes are order based upon log2-fold change in expression.

| Ensemble ID | Gene Name | Description | log2 Fold Change | p-value | adjusted p-value |
| --- | --- | --- | --- | --- | --- |
| ENSMUSG00000005892 | *Trh* | thyrotropin releasing hormone | 2.879761 | 1.02E-06 | 7.81E-04 |
| ENSMUSG000000060131 | *Atp8b4* | ATPase, class I, type 8B, member 4 | 2.076196 | 4.16E-06 | 2.57E-03 |
| ENSMUSG000000069372 | *Ctxn3* | cortexin 3 | 1.554173 | 6.52E-06 | 3.46E-03 |
| ENSMUSG000000062609 | *Kcnj15* | potassium inwardly-rectifying channel, subfamily J, member 15 | 1.447912 | 7.43E-05 | 2.37E-02 |
| ENSMUSG000000045053 | *Kcng3* | potassium voltage-gated channel, subfamily G, member 3 | 1.433555 | 2.70E-19 | 7.38E-16 |
| ENSMUSG000000031603 | *Fgf20* | fibroblast growth factor 20 | 0.868887 | 7.82E-05 | 2.40E-02 |
| n.a. | *Gm30575* | predicted gene, 30575 | 0.822134 | 1.40E-05 | 5.94E-03 |
| ENSMUSG000000047642 | *D930020B18Rik* | RIKEN cDNA D930020B18 gene | 0.745807 | 4.75E-06 | 2.67E-03 |
| ENSMUSG000000115012 | *9630009A06Rik* | RIKEN cDNA 9630009A06 gene | 0.743960 | 7.92E-05 | 2.40E-02 |
| n.a. | *Gm38650* | predicted gene, 38650 | 0.724578 | 1.31E-05 | 5.71E-03 |
| n.a. | *Gm46899* | predicted gene, 46899 | 0.656898 | 1.38E-06 | 1.01E-03 |
| ENSMUSG000000089798 | *1700028K03Rik* | RIKEN cDNA 1700028K03 gene | 0.644188 | 1.32E-04 | 3.40E-02 |
| ENSMUSG000000085525 | *Gm13166* | Predicted gene 13166 | 0.632088 | 1.00E-09 | 1.60E-06 |
| ENSMUSG000000019989 | *Enpp3* | ectonucleotide pyrophosphatase/phosphodiesterase 3 | 0.622093 | 4.49E-09 | 5.73E-06 |
| ENSMUSG000000020212 | *Mdm1* | transformed mouse 3T3 cell double minute 1 | 0.621684 | 7.49E-22 | 2.86E-18 |
| ENSMUSG000000087095 | *Emx2os* | Emx2 opposite strand/antisense transcript (non-protein coding) | 0.613505 | 5.18E-06 | 2.83E-03 |
| n.a. | *Gm52889* | predicted gene, 52889 | 0.601715 | 1.04E-04 | 2.89E-02 |
| n.a. | *Gm53737* | predicted gene, 53737 | 0.590884 | 3.68E-05 | 1.28E-02 |
| n.a. | *Gm52704* | predicted gene, 52704 | 0.543151 | 1.18E-04 | 3.18E-02 |
| ENSMUSG000000021209 | *Ppp4r4* | protein phosphatase 4, regulatory subunit 4 | 0.526342 | 2.17E-06 | 1.54E-03 |
| n.a. | *Gm54130* | predicted gene, 54130 | 0.526132 | 2.14E-04 | 4.75E-02 |
| ENSMUSG000000113379 | *Gm46436* | Predicted gene, 46436 | 0.521977 | 9.81E-05 | 2.80E-02 |
| ENSMUSG000000027397 | *Slc20a1* | solute carrier family 20, member 1 | 0.510199 | 2.22E-09 | 3.26E-06 |
| n.a. | *Gm38485* | predicted gene, 38485 | 0.508928 | 3.00E-05 | 1.06E-02 |
| ENSMUSG000000021281 | *Tnfaip2* | tumor necrosis factor, alpha-induced protein 2 | 0.507720 | 1.51E-04 | 3.84E-02 |
| ENSMUSG000000023467 | *Tulp2* | tubby-like protein 2 | 0.504044 | 1.56E-04 | 3.86E-02 |
| ENSMUSG000000097187 | *Gm19426* | predicted gene, 19426 | 0.498688 | 1.23E-04 | 3.26E-02 |
| ENSMUSG000000046169 | *Adamts6* | a disintegrin-like and metallopeptidase (reprolysin type) with thrombospondin type 1 motif, 6 | 0.493700 | 3.57E-07 | 2.97E-04 |
| n.a. | *Gm53541* | predicted gene, 53541 | 0.420842 | 1.91E-04 | 4.46E-02 |
| ENSMUSG000000019789 | *Hey2* | hairy/enhancer-of-split related with YRPW motif 2 | 0.416916 | 8.77E-07 | 6.98E-04 |
| ENSMUSG000000057060 | *Slc35f3* | solute carrier family 35, member F3 | 0.409773 | 2.61E-05 | 9.66E-03 |
| ENSMUSG000000037492 | *Zmat4* | zinc finger, matrin type 4 | 0.370916 | 1.18E-04 | 3.18E-02 |
| ENSMUSG000000031736 | *Crnde* | colorectal neoplasia differentially expressed (non-protein coding) | 0.362081 | 1.67E-05 | 6.65E-03 |
| ENSMUSG000000053332 | *Gas5* | growth arrest specific 5 | 0.346093 | 1.94E-04 | 4.46E-02 |
| ENSMUSG000000010175 | *Prox1* | prospero homeobox 1 | 0.319460 | 2.90E-08 | 2.77E-05 |
| ENSMUSG000000039191 | *Rbpj* | Recombination signal binding protein for immunoglobulin kappa J region | 0.263601 | 2.54E-05 | 9.66E-03 |

**Table 3:**
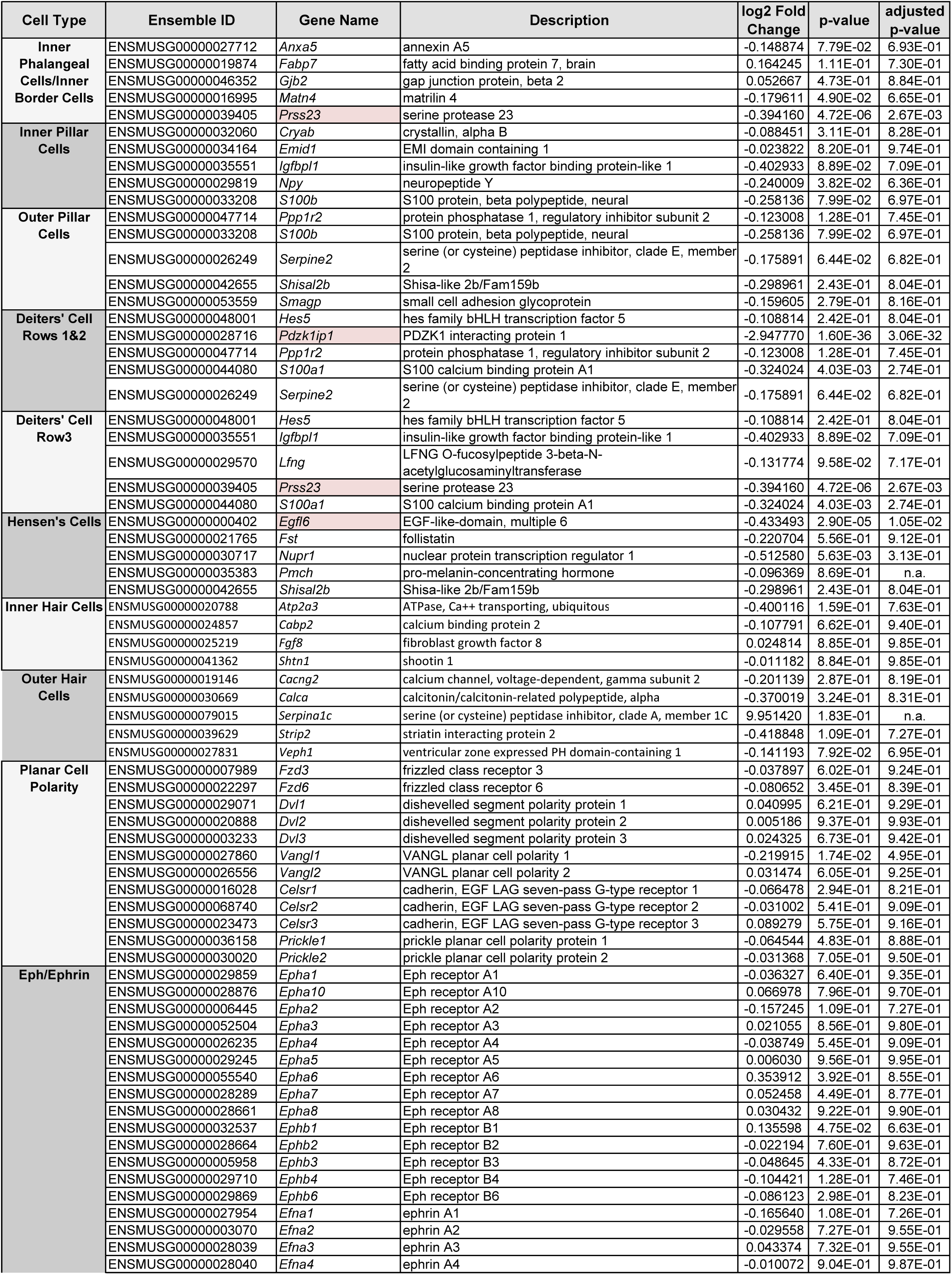
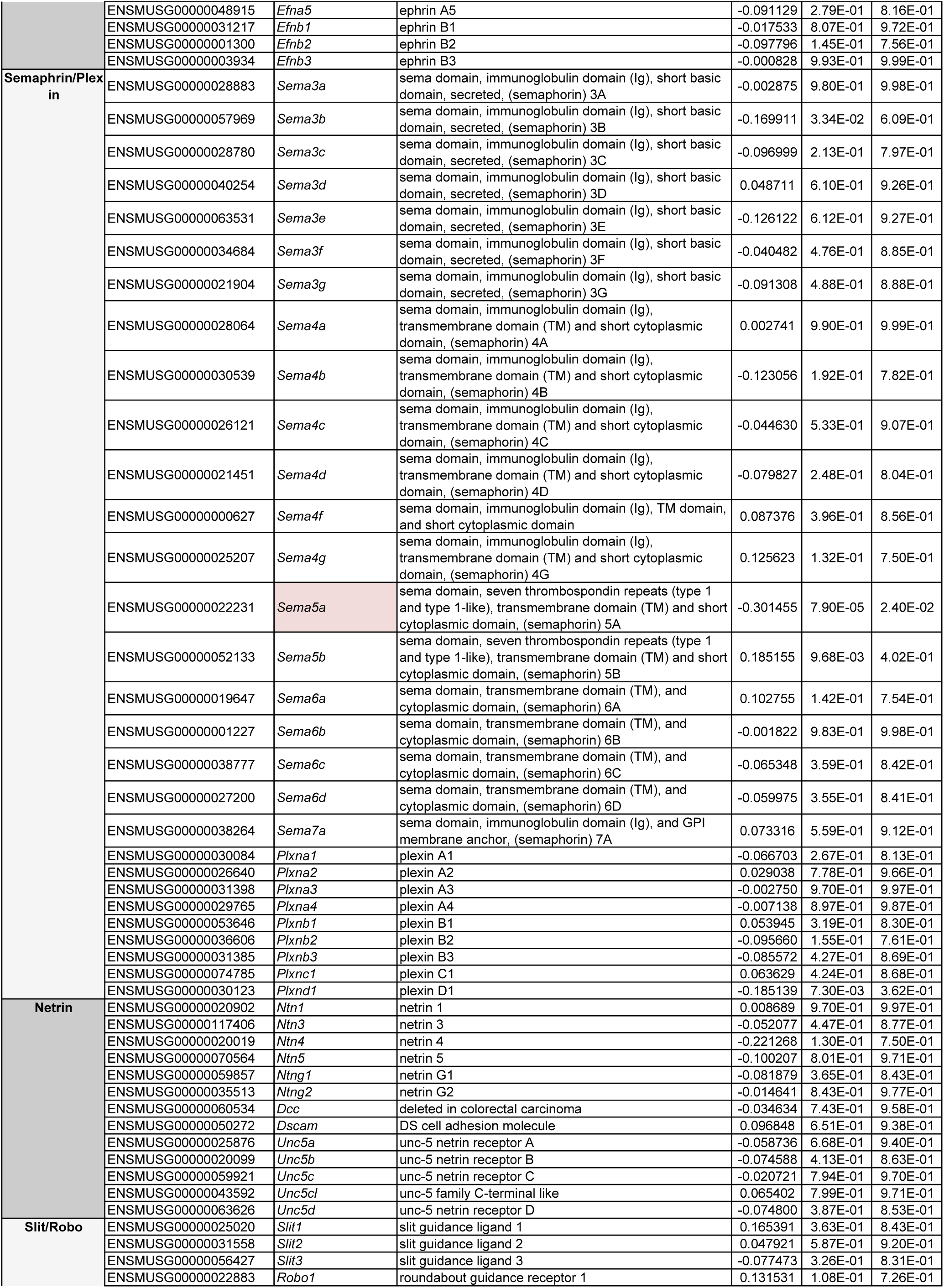
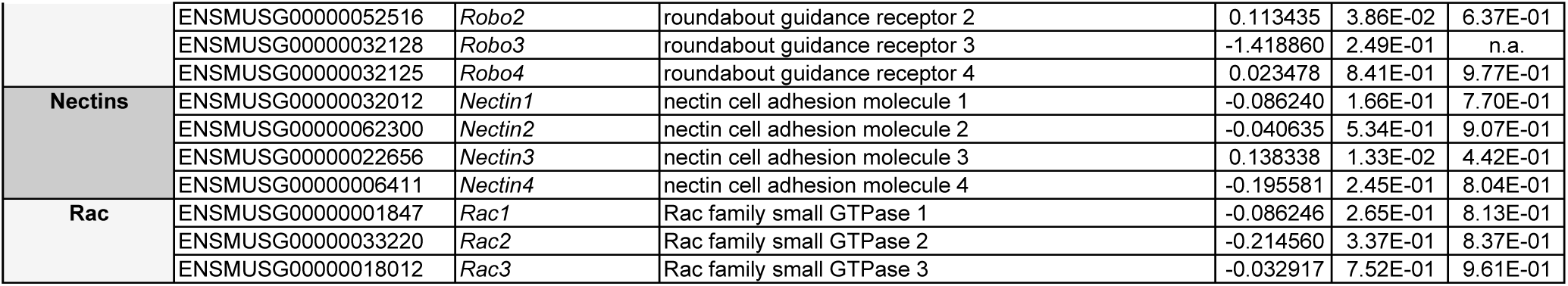
Gene Family and Cell Type Signature Gene Expression Changes in *Emx2*-Cre; *Prox1* cKO. List of select genes and the measured changes in their expression between *Emx2*-Cre; *Prox1* cKOs and littermate controls. Cell type signatures are sets of 4-5 genes whose expression distinguished that particular cell type from other cells in the P1 mouse organ of Corti based upon scRNAseq datasets (Kolla, Kelly et al. 2020). Also listed are genes that might influence type II SGN innervation of the lateral compartment. These include all classical axon guidance ligands and their receptors and PCP genes, as well as Nectins 1-3 and Rac1-3. Red shading indicates genes with significant changes in the cKO.

Despite the Type II SGN axon guidance phenotype seen in the *Emx2*-Cre; *Prox1* cKO, no PCP genes were impacted by the loss of *Prox1* (Fig.5D, Supp. Tables 1-3). This included *Vangl2* and the non-canonical Wnt/PCP receptors *Frizzled3* and *Frizzled6* that have been shown to direct peripheral axon turning (Ghimire, Ratzan et al. 2018, Ghimire and Deans 2019). *Rac1* and *Nectin3* levels were also not changed despite their contribution to axon guidance in the mouse cochlea (Clancy, Xie et al. 2025). The only axon guidance gene impacted was the Semaphorin ligand *Sema5a* which showed a 0.30 log2fold decrease in the *Emx2*-Cre; *Prox1* cKO (Supp. Table1). *Sema5a* is expressed in the otic mesenchyme surrounding the developing cochlea at this stage (Rose, Manilla et al. 2023) and it is possible the mesenchymal cells were present in tissue used for RNA isolation due to the mechanical nature of organ of Corti dissection. However, when queried using the gene Expression Analysis Resource (gEAR) (Orvis, Gottfried et al. 2021), *Sema5a* could also be found in small numbers of IPCs, OPCs and Deiters’ cells of the mouse organ of Corti at P1 and therefore may also be present earlier during innervation of the lateral compartment.

The RNAseq data also did not suggest that there well cell fate changes in the *Emx2*-Cre; *Prox1* cKO. Previous scRNAseq characterization of the organ of Corti identified sets of genes whose co-expression signified supporting cell type identity (Kolla, Kelly et al. 2020). When the expression of these genes was compared between *Emx2*-Cre; *Prox1* cKOs and littermate controls there were no differences that would indicate changes in supporting cell identity after *Prox1* gene deletion (Fig.5E). Three genes were downregulated significantly, but they were not expressed by the same cell type. Moreover, there was not a corresponding increase in expression for genes that characterized other cell types as would be expected following a cell fate transition in the absence of *Prox1*. Similarly, genes distinguishing IHCs from OHCs were not impacted (Supp. Table3). Gene Ontology (GO) analysis of down-regulated genes using Metascape (Zhou, Zhou et al. 2019) also did not identify changes in PCP or axon guidance pathways (Fig.5F). Instead, the most impacted GO enriched terms related to the positive regulation of cell migration, and locomotion was the most impacted GO biological process.

#### Axon guidance errors in the *Prox1* cKO are a secondary phenotype

In other systems, SEMA5a is an axon guidance cue that acts as either an attractant or a repellent depending upon developmental context (Kantor, Chivatakarn et al. 2004). If expressed by lateral compartment supporting cells SEMA5a could act as a guidance cue directing type II peripheral turning towards the cochlear base. However, in a *Sema5a* KO (Matsuoka, Chivatakarn et al. 2011) peripheral axons labeled for Neurofilament and evaluated at P0 were indistinguishable from littermate controls (Fig.6A). Therefore, decreased *Sema5a* expression is not expected to cause the innervation errors that occur in *Emx2*-Cre; *Prox1* CKOs.

**Figure 6:**
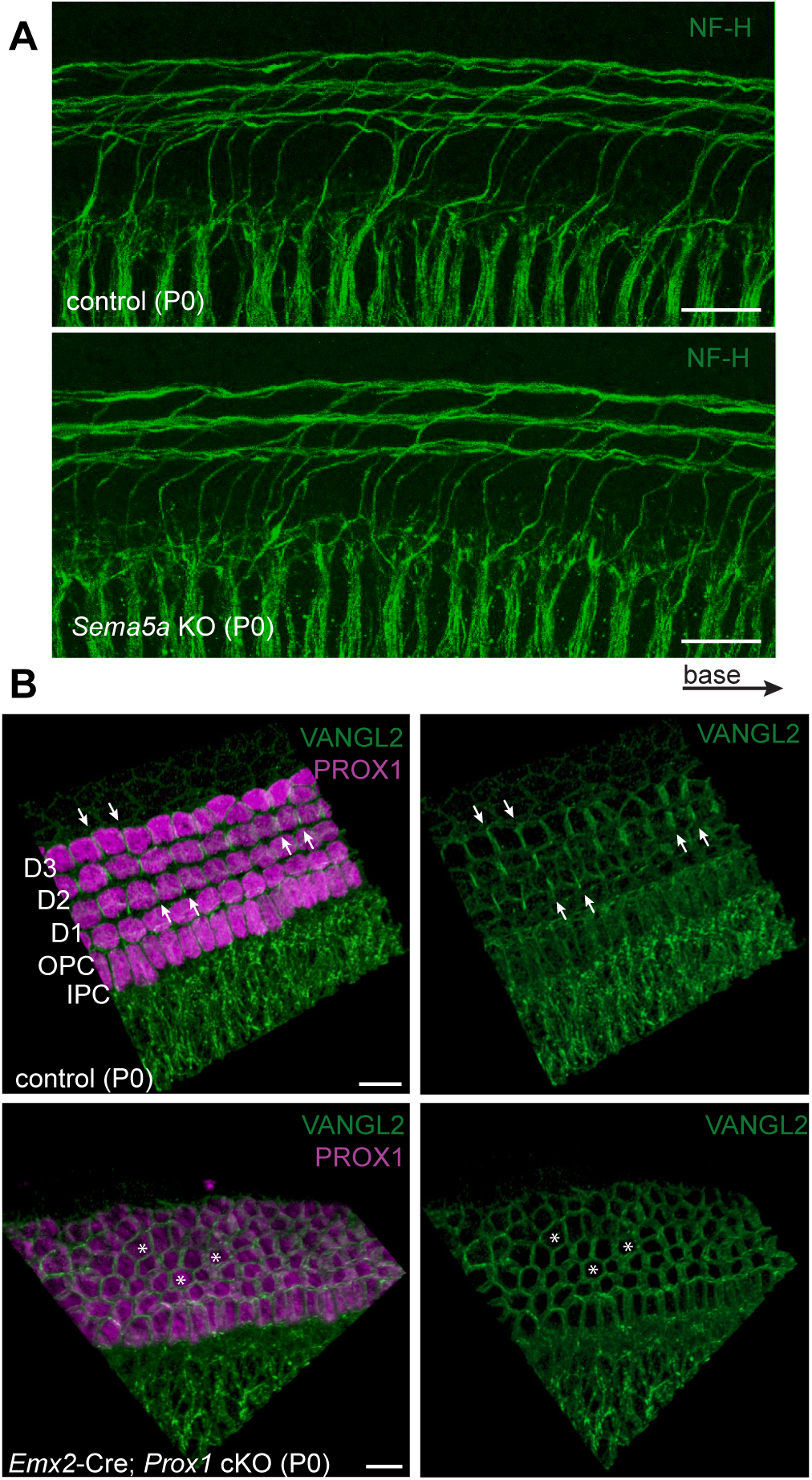
The polarized distribution of VANGL2 is lost *Emx2-*Cre; *Prox1* cKOs. (**A**) Evaluation of peripheral axon turning in *Sema5a* KO mice compared to littermate controls at P0 by wholemount immunolabeling against NF-H. (**B**) Immunofluorescent images of VANGL2 wholemount organ of Corti viewed as 3D-reconstructions. Arrowheads mark examples of polarized enrichment of Vangl2 at the intercellular junctions between neighboring supporting cells in littermate controls (n=4) while asterisks mark examples of cells lacking this polarity in *Emx2-*Cre; *Prox1* cKOs (n=3) at P0. Prox1 immunolabeling marks cell nuclei. Images are collected from the basal turn of the cochlea. Scale bars are 20µm.

In the absence of transcriptional changes, the axon guidance defects seen in the *Emx2-*Cre; *Prox1* cKOs may be secondary to the disorganization of supporting cells within the environment that the growth cone traverses. This may be especially relevant for the PCP proteins which must be asymmetrically distributed to mediate intercellular signaling in the ear (Sienknecht, Anderson et al. 2011, Stoller, Roman et al. 2018). Moreover, in the area where Type II SGN growth cones turn, VANGL2 is enriched at the intercellular junctions between neighboring IPCs, OPCs and Deiters’ cells where it has been proposed to act as a guidance cue (Ghimire, Ratzan et al. 2018). This distribution is readily detected by immunofluorescent labeling of VANGL2 shortly after axon turning is complete at P0 (Fig.6B). Notably, where supporting cells are disorganized in the *Emx2*-Cre; *Prox1* cKO, the polarized distribution of VANGL2 is also lost. In the cKO, VANGL2 protein is more likely to surround individual cells than to be distributed in an organized pattern that might provide spatially restricted guidance cues (Fig.6B).

#### *Prox1* contributes to the differentiation of lateral compartment supporting cells

The continued expression of gene clusters that define lateral compartment supporting cell types suggests that their identity is correctly specified in the absence PROX1. These cells also continue to transcribe *Prox1* mRNA which further indicates that they are developing as lateral compartment supporting cells. To determine whether PROX1 regulates cell differentiation instead of specification, markers expressed at later stages of development and therefore not captured by RNAseq at P0, were evaluated by immunofluorescent labeling. One such marker is the cell surface protein CD44 which can be detected at P2 and distinguishes the OPCs from IPCs (Fig.7A) (Hertzano, Puligilla et al. 2010). In littermate controls, CD44 labels a single row of PROX1-expressing cells that are the OPCs in addition to the Claudius cells and greater epithelial ridge cells that flank the organ of Corti. However, in *Emx2*-Cre; *Prox1* cKOs this marker of OPC differentiation is lost from cells expressing mutant PROX1. Another characteristic of supporting cell differentiation is the down regulation of PROX1 expression which normally becomes restricted to OPCs and third row Deiters’ cells at P7. This downregulation does not occur in the cKO where mutant PROX1 expression is retained in all lateral compartment supporting cells and the IPCs (Fig.2A). Despite this, IPC differentiation may not be impacted because their apical processes are immunolabeled by the marker p75/NTR (Mueller, Jacques et al. 2002) and can be seen as a strip positioned between the IHCs and first row of OHCs at the level of the reticular lamina (Fig. 7C). IPCs also develop a distinctive oblong nucleus positioned at the nuclear layer that is not impacted by the loss of *Prox1* (Fig.7B). For reference the imaging planes for the reticular lamina and supporting cell nuclear layers are annotated in Fig.1A.

**Figure 7:**
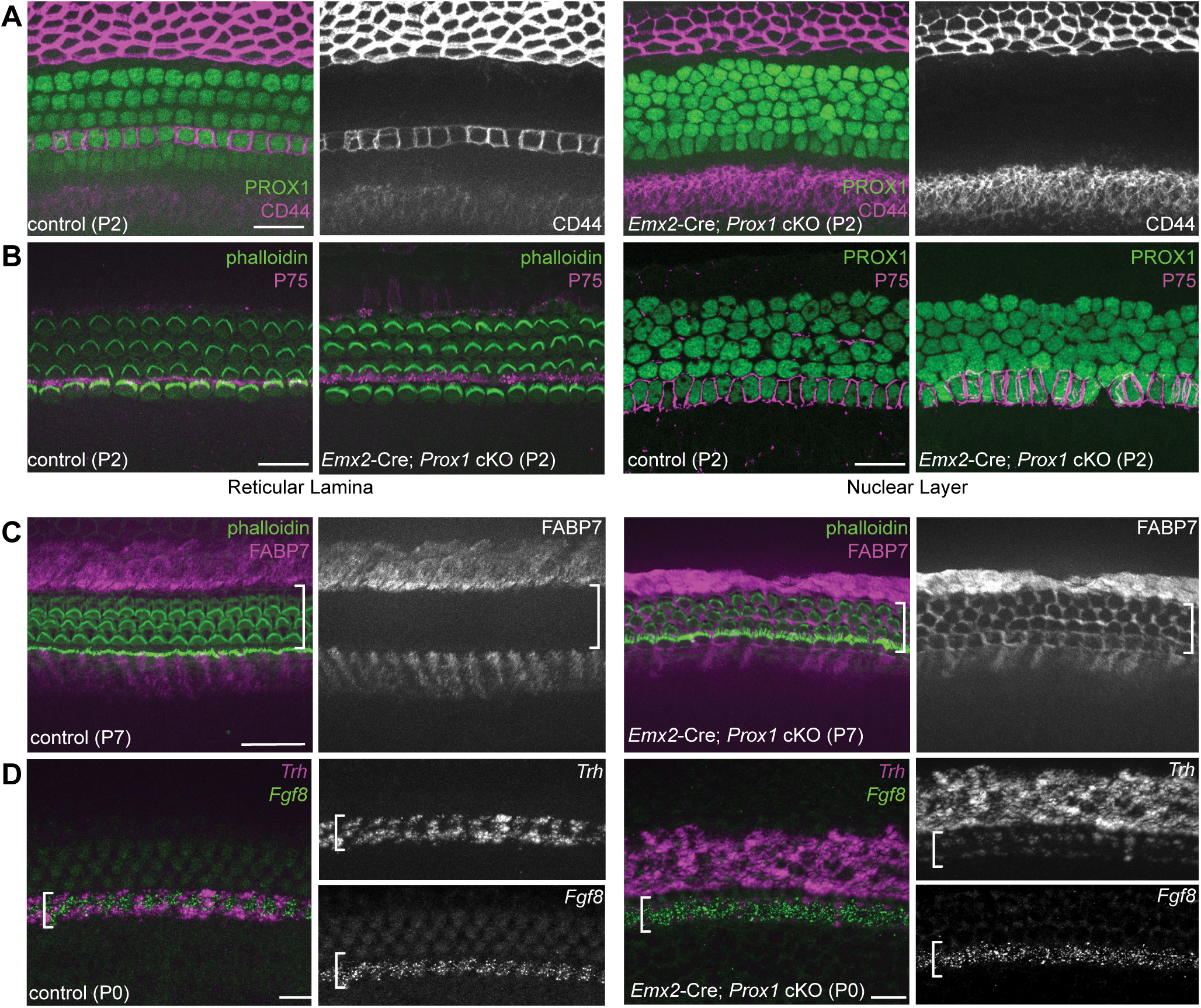
Lateral compartment supporting cell differentiation appears incomplete in *Emx2-*Cre; *Prox1* cKOs. (**A**) Wholemount immunofluorescence images of PROX1 and the OPC protein CD44 in the developing organ of Corti shows the absence of CD44 expression in *Emx2-*Cre; *Prox1* cKOs (n=3) compared to littermate controls (n=2) when evaluated at P2. (**B**) Immunolabeling for the IPC protein P75-NTR shows no changes in the apical projections of IPCs visualized at the reticular lamina or the number or distribution of IPCs at the nuclear layer. (**C**) Immunolabeling for the medial compartment supporting cell protein FABP7 shows an expansion of FABP7 into the lateral compartment in *Emx2-*Cre; *Prox1* cKOs (n=3) compared to littermate controls (n=2) at P7. Bracket marks the boundaries of the lateral compartment based on the distribution of Phalloidin-stained OHCs. (**E**) fISH showing the distribution of the medial gene *Trh* relative to IHCs expressing *fgf8* and expansion of *Trh* expression in *Emx2-*Cre; *Prox1* cKOs (n=4) compared to littermate controls (n=5) at P0. Brackets mark the position of the IHC row. Scale bars are 20µm.

Lateral compartment supporting cells can also be distinguished from medial compartment cells by markers that they don’t express including FABP7. At P7, FABP7 normal labels inner phalangeal cells in the medial compartment and Hensen’s cells that are located on the opposite side of the lateral compartment. However, in *Emx2*-Cre; *Prox1* cKOs at P7, FABP7 immunolabeling expands to include cells throughout the lateral compartment (Fig.7C). Another inner phalangeal marker is *thyrotropin releasing hormone* (*Trh*) which is also the DEG that exhibited the greatest increased expression in the *Emx2*-Cre; *Prox1* cKO at P0 (Fig.5A). Evaluation by fluorescent ISH shows that this upregulation occurs because, like FABP7, *Trh* expression expands to include the entire lateral compartment of the organ of Corti. Together these observations suggest that PROX1 promotes lateral supporting cell differentiation by promoting expression of genes enriched in those cells while also preventing the expression of genes normally found in the medial compartment.

Cellular differentiation can also be unambiguously established based upon morphological criteria, and a distinctive structural outcome of pillar cell differentiation is formation of the tunnel of Corti. The tunnel forms between IPCs and OPCs during the course of their differentiation and is the anatomical feature separating medial and the lateral compartments after P10 (Sher 1971). Tunnel of Corti formation was evaluated in *Emx2*-Cre; *Prox1* cKOs by immunofluorescent labeling of cryosectioned tissue at P14. DAPI staining was used to identify supporting cell and hair cell nuclei, while tissue auto-fluorescence revealed tissue boundaries and highlighted the tunnel of Corti (Fig.8A). In the cKO cochlea, the tunnel fails to develop in the apical turns and is incompletely developed in the middle turns but appears normal in the cochlear base (Fig.8B). Another striking example of morphological differentiation is the extension of the phalangeal process from Deiters’ cells which projects from their cell body located beneath one OHC towards the reticular lamina where it inserts between the apical surfaces of two neighboring OHCs. The phalangeal process can be readily detected by immunolabeling its primary structural component Tubulin. In the *Emx2*-Cre; *Prox1* cKO the differentiation of the phalangeal process is disrupted and it frequently projects alongside same OHC contacting the Deiters’ cell soma rather than projecting towards the cochlear apex Fig.8B). Altogether these aspects of the cKO phenotype demonstrate that PROX1 is required for supporting cell differentiation and that in its absence there are structural changes that are likely to impact auditory function.

**Figure 8:**
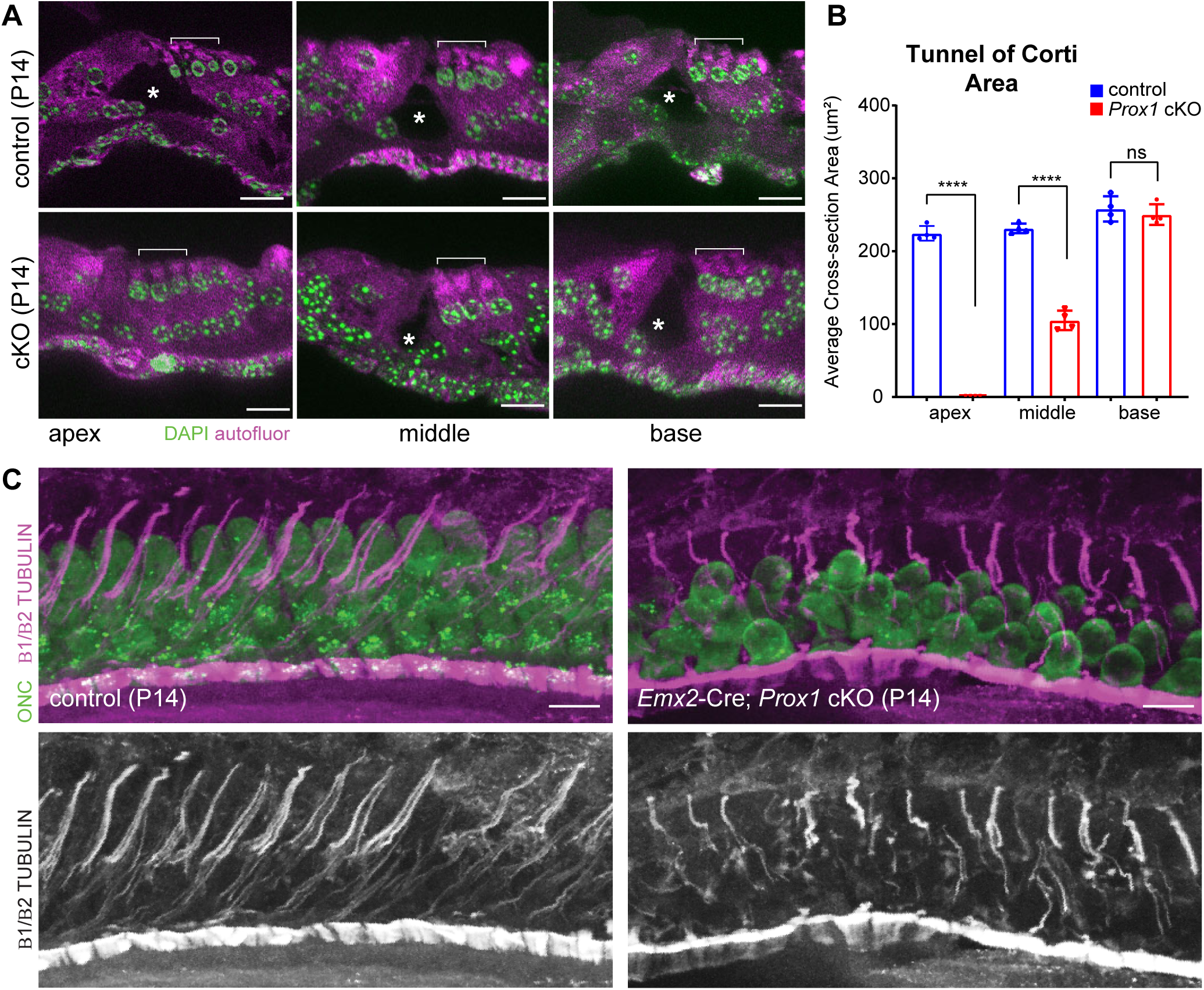
Structural changes in lateral compartment supporting cells. (**A**) Cochlear sections cut from P14 *Emx2*-Cre; *Prox1* cKOs and littermate controls labeled with the nuclear dye DAPI and imaged against background autofluorescence from a second channel. Asterisks mark the tunnel of Corti center and brackets the position of the three rows of OHC nuclei. (**B**) Area of the tunnel of Corti in cochlear sections for *Emx2-*Cre; *Prox1* cKOs (n=4) and littermate controls (n=4) at 3 positions along the length of P14 cochleae. Error bars show standard deviation, and asterisks indicate statistical significance for differences between genotypes using Student’s *t*-test (***p<0.0001). (**C**) Deiters’ cell phalangeal processes immunolabeled with antibodies against β1β2 Tubulin reveals differences in their structural morphology and organization relative to OHCs in *Emx2-*Cre; *Prox1* cKOs (n=3) compared to littermate controls (n=3) at P14. OHCs are labeled using antibodies against Oncomodulin. Scale bars are 10µm (A).

## Discussion

In this study we used a *Prox1* cKO mouse line to investigate the role of the PROX1 transcription factor during mouse organ of Corti development. The mammalian organ of Corti is divided into functional domains, with the lateral compartment containing the OHCs and associated supporting cells that enable cochlear amplification. We hypothesized that *Prox1* would play an important role in lateral compartment development because it is one of the first transcription factors to be expressed in this domain and is restricted to IPCs, OPCs and Deiters’ cells. Consistent with this rationale we observed an increase in supporting cell numbers in the *Emx2*-Cre; *Prox1* cKO and as a result, disorganized rows of supporting cell nuclei and apical projections to the reticular lamina. We also found that proper innervation of the lateral compartment is dependent upon PROX1 functions in supporting cells rather than the innervating neurons, and that the axon guidance errors we observed were not due to changes in axon guidance gene expression. Instead, we propose that growth cone turning errors are secondary to supporting cell disorganization. Finally, by comparing RNAseq data with a publicly available scRNAseq dataset we conclude that the primary function of Prox1 is regulating supporting cell differentiation rather than the specification of supporting cell types. It is noteworthy that disrupted differentiation resulted in pronounced structural changes in *Prox1* cKOs, including the incomplete formation of the tunnel of Corti and altered Deiters’ cell projections towards the reticular lamina, that are likely to disrupt auditory function.

### Prox1 Regulates supporting cell Patterning and Number

Our findings demonstrate that PROX1 function contributes to the precise cell patterning that characterizes hair cells and supporting cells in the mammalian organ of Corti. The modest ∼24% increase in supporting cells does not result in an additional row of supporting cell nuclei that would be expected if proper patterning was maintained. Instead supporting cell nuclei are misplaced in *Prox1* cKOs and their orderly organization is lost. These supernumerary supporting cells are only found in the lateral compartment, consist of cells that continue to express mutant PROX1, and are likely to be both OPCs and Deiters’ cells based upon the distribution of their phalangeal projections to the reticular lamina. In contrast, OHC number is not changed in *Emx2*-Cre; *Prox1* cKOs. This is striking because hair cell and supporting cell numbers are usually tightly coordinated, and in the converse situation where extra hair cells are produced there is a corresponding increase in supporting cell number (Hayashi, Cunningham et al. 2007).

In the embryonic retina, the loss of *Prox1* prevents the development of horizontal cells and an additional round of progenitor cell division leads to increases in other retinal cell types (Dyer, Livesey et al. 2003). However, an important difference between the retina and cochlea is that retinal progenitors continue to divide throughout development whereas in the cochlea sensory progenitors exit the cell cycle at the same time. Moreover, we did not find evidence for either increased cell division or decreased apoptosis in *Emx2*-Cre; *Prox1* cKO cochleae. An alternative explanation is that the extra supporting cells are derived from a second population of cells that transform into supporting cells between E17.5 and P0 and then express mutant PROX1. This could occur if Hensen’s cells were transformed into Deiters’ cells except that corresponding reductions in Hensen’s cells or other cell types were not observed in the *Emx2*-Cre; *Prox1* cKO. Alternatively neural crest cells that are normally migrating towards the outer sulcus at this stage could be recruited into the organ of Corti and commit to a PROX1-expressing cell fate. Interestingly, both scenarios would require that the organ of Corti has a census mechanism that counts OPC and Deiters’ cell number, and a compensatory mechanism that can recruit additional cells if those counts are low.

### Prox1 in Type II SGN Pathfinding

The OHCs in the lateral compartment are exclusively innervated by type II SGNs which extend a peripheral axon into this compartment that makes a characteristic 90 degree turn towards the base of the cochlear spiral. Turning is regulated by guidance cues expressed in lateral compartment supporting cells. For example, PCP signaling directs axon turning, and Vangl2 and Fzd3 are asymmetrically distributed at intercellular junctions between neighboring supporting cells (Ghimire, Ratzan et al. 2018, Ghimire and Deans 2019). The cell adhesion molecule Nectin3 and the actin cytoskeletal regulator Rac1 also act non-autonomously from the lateral compartment supporting cells to direct turning (Clancy, Xie et al. 2025). Since PROX1 is expressed in both lateral compartment supporting cells and the SGN it could regulate the transcription of factors acting on one or both sides of this axon guidance signal. Previously it was proposed that *Prox1* was required in the type II SGN to direct axon turning and OHC innervation based on cKOs generated using *Nestin*-Cre (Fritzsch, Dillard et al. 2010). However, when we targeted *Prox1* in the SGN using the more specific *Bhlhe22*-Cre line there were no turning errors. One explanation might be differences in levels of Cre expression between these two lines or the efficiency of Cre-mediated recombination in different SGN sub-types. However, since *Bhlhe22*-Cre is active early in development (Appler, Lu et al. 2013) and we see extensive deletion of the targeted *Prox1* exons by P0 (Fig.4A) we have to conclude that this line is sufficient to generate the intended SGN-restricted cKO. Thus, while our data do not rule out important functions for PROX1 in the SGN, it is not consistent with an autonomous role for PROX1 in the type II SGN during peripheral axon turning.

In contrast, pronounced turning errors occurred when *Prox1* was selectively removed from the organ of Corti using *Emx2*-Cre. This non-autonomous function for PROX1 highlights the important role that supporting cells play in guiding neuronal development in the cochlea. Candidate factors that might regulate axon turning were evaluated in the *Emx2*-Cre; *Prox1* cKO by quantification using RNAseq with the expectation that essential factors would be down-regulated. However, there were no changes in PCP gene expression, axon guidance cue or receptor expression, or the levels of *Nectin3* and *Rac1* mRNA. GO analysis also did not identify changes in axon guidance or related development pathways. The greatest changes were for genes associated with cell migration or locomotion and these are more likely to be related to the cellular disorganization in the *Prox1* cKO than movements of the growth cone. Based upon these observations we propose that PROX1 does not regulate peripheral axon turning directly, and that instead the turning errors are secondary to the disorganization of supporting cells. Consistent with this interpretation, the polarized distribution of the core PCP protein VANGL2 at the junctions between neighboring supporting cells is lost in the *Emx2*-Cre; *Prox1* cKO (Fig.6). A plausible explanation is that changes in the distribution of PCP proteins themselves, or proteins whose distribution might be regulated by VANGL2 including Rac1 and NECTIN3 (Clancy, Xie et al. 2025) lead to the axon turning errors we see in *Prox1* cKOs.

### Cell type specification versus differentiation

Altogether these data demonstrate that PROX1 is necessary for the proper differentiation of lateral compartment supporting cells but not their specification to become OPCs or Deiters’ cells. The earliest and most salient evidence for proper lateral fate specification in the *Emx2*-Cre; *Prox1* cKO is the persistent expression from the *prox1* gene promoter revealed by immunolabeling for mutant PROX1 protein. If these cells had not been specified as lateral compartment supporting cells they would not continue to express PROX1. However, the expression of a single marker is not sufficient to specify cell identity, and single-cell transcriptomics have shown that identity is better defined by larger transcriptional signatures. By comparing the levels of expression for cell-type specific RNA signatures identified by scRNAseq (Kolla, Kelly et al. 2020) to our whole-tissue RNAseq of *Emx2*-Cre; *Prox1* cKOs, we found no evidence indicating changes in the specification or the relative distribution of supporting cell types (Fig.5 and Supp. Tables 1&2). A limitation of this approach may be that signature genes can be expressed in multiple cell types, and the absence of one cell type might not produce a great enough change in gene expression to be detected by whole tissue RNAseq. However, this seems unlikely since the approach was sensitive enough to detect an increase in *Prox1* transcripts (Fig.5C) resulting from the increase in lateral compartment supporting cells in the cKO. It is also noteworthy that lateral compartment cells upregulate the expression of genes that are normally found in the medial compartment such as *Trh* and *FABP7*. This may indicate that lateral compartment specification is incomplete and that in the absence of *Prox1* cells have a mixed identity. Alternatively, one aspect of supporting cell differentiation may be the active repression of genes found in other cell types. Either scenario is consistent with the persistent transcription from the *Prox1* allele that we see in the *Emx2*-Cre; *Prox1* cKO.

Separate evidence that PROX1 is necessary for supporting cell differentiation comes from the changes in supporting cell structures that are formed at later stages of development. One example is opening of the tunnel of Corti between the IPCs and OPCs. In the *Prox1* cKO the tunnel fails to form in the apical turns of the cochlea and is narrower in the middle turns. The graded progression of this phenotype is consistent with PROX1 regulating differentiation and not specification since a cochlea without OPCs should be missing the tunnel along its entire length. A second example are the distinctive phalangeal processes that emanate from Deiters’ cells and project towards the apical turn of the cochlea. These processes are important for the structural integrity of the organ of Corti and develop postnatally and after Deiters’ cell specification in the embryo. We find that in the absence of PROX1 phalangeal process extension is stunted so that they project directly towards the reticular lamina rather than tilting towards the cochlear apex. Altogether the phenotypes we observed in *Prox1* cKO mice demonstrate that PROX1 plays an important role in organ of Corti development by regulating OPC and Deiters’ cell differentiation and number.

## Acknowledgements

Research reported in this publication was supported by the National Institute On Deafness And Other Communication Disorders of the National Institutes of Health under Award Number R01DC018040. The content is solely the responsibility of the authors and does not necessarily represent the official views of the National Institutes of Health.

## Materials and Methods

### Mouse strains and husbandry

Mice housed at the University of Utah were maintained in accordance with Institutional Animal Care and Use Committee (IACUC) protocol #1498. *Sema5a* KO mice were maintained at Georgetown University in accordance with IACUC protocol #1147. Individual lines at the Univ. of Utah were maintained by heterozygous backcross to B6129SF1/J hybrid females (Jax strain#101043) and mice with the *Prox1* ^LoxP/LoxP^ genotype were maintained by homozygous intercross. *Sema5a* mice were maintained by backcross to C57Bl/6N mice. All genotyping was performed using allele-specific PCR genotyping reactions (primer sequences available upon request). For timed breeding and tissue collection, noon on the day a vaginal plug was observed was designated as embryonic day 0.5 (E0.5), and the day of birth postnatal day 0 (P0). Conditional knockouts were generated by crossing *Emx2* ^Cre/wt^; *Prox1* ^LoxP/wt^ males or *Bhlhe22* ^Cre/wt^; *Prox1* ^LoxP/wt^ males with *Prox1 ^LoxP/LoxP^* females. Only Cre-positive progeny were analyzed and embryos or pups of both sexes were used for experimentation. *Sema5a* KO mice were generated by crossing *Sema5a +/-* males and *Sema5a ^+/-^* females*. Emx2* Cre mice (Kimura, Suda et al. 2005) were provided by S.Aizawa (Riken Institute), *Bhlhe22*-Cre mice (Ross, Mardinly et al. 2010) were provided by M.Greenberg (Harvard Medical School), *Prox1* cKO mice (Harvey, Srinivasan et al. 2005) were provided by G.Oliver (Northwestern University), and *Sema5a* mice (Matsuoka, Chivatakarn et al. 2011)were provided by A.Kolodkin (Johns Hopkins University).

### In-Situ Hybridization

Cochleae were prepared for ISH by fixing temporal bones overnight at 4°C using 4% paraformaldehyde (PFA) in 67 mM Sorensons’ phosphate buffer (pH 7.4) followed by storage at -20°C in 100% MeOH as needed. Prior to hybridization, tissue was dissected to expose the sensory epithelia of the cochlea, rehydrated through a decreasing MeOH gradient and transferred to 1XPBS with 0.05% Tween-20 (PBSt). Dissected tissue was treated with Proteinase K (30ug/mL, Ambion #Am2546) for 12 minutes at room temperature (RT), rinsed with PBSt, and post-fixed at RT using 4% PFA/Sorensons’ buffer for 20 mins. Fluorescent *in-situ* hybridizations were performed in 96-well plates using the HCR protocol (Choi, Schwarzkopf et al. 2018). In brief, hybridization buffer contained 16nm of individual mRNA probes and hybridization occurred overnight at 37°C in a humidified container. Hairpin amplification used 4µM Metastable DNA Hairpins and samples were incubated overnight at room temperature. Fluorescent images were captured by structured illumination microscopy using a Zeiss Axio Imager M.2 with ApoTome.2 attachments and an Axiocam 506 monochrome camera. For histochemical *in situ* hybridization tissue was blocked, permeabilized, and hybridized with a digoxigenin-labeled anti-sense probe that was detected using alkaline phosphatase conjugated anti-digoxigenin antibodies (Roche). Following the alkaline phosphatase histochemistry, tissue was imaged using the Zeiss Axio Imager M.2 and Axiocam 105 color camera. Detailed ISH protocols for fluorescent and conventional ISH are available upon request. HCR probes used in this study recognized the *Fgf8*, *Gfi-1* and *Trh* mRNAs and the *Prox1* probe was specific for the exons targeted in *Prox1* cKOs (Custom probe sets from Molecular Instruments). For digoxigenin-labeled probes, a *Prox1* template was generated by PCR amplification of whole embryo cDNA using a primer pair flanking the targeted exons (5’-CACCCGCTACCCCAGCTCCAACAT-3’, 5’-TTTCAATGCCATCATCGCGGGC-3’). The amplified cDNA was TOPO-cloned into a plasmid containing T7 and SP6 promoters to facilitate RNA probe transcription.

### Immunofluorescent labeling

Brain tissue was removed from bisected heads and temporal bones fixed overnight at 4°C using 4% paraformaldehyde (PFA) in 67 mM Sorensons’ phosphate buffer (pH 7.4). For wholemount labeling, cochleae were micro-dissected and permeabilized for 30 minutes at RT using blocking solution (5% normal donkey serum, 1% BSA, PBS) supplemented with 0.5% Triton-X100. Primary antibodies and phalloidin AlexaFluor 488 (Invitrogen, A12379; 1:1000) were diluted in blocking solution supplemented with 0.1% Tween-20 and incubated overnight at 4°C. Following incubation, tissues were washed three times in PBS with 0.05% Tween-20, and incubated overnight at 4°C with species-specific Alexa-conjugated secondary antibodies diluted in blocking solution with 0.1%Tween-20 (catalog numbers TBD, diluted 1:1500). After a final round of washing, tissues were mounted in Prolong Gold (Molecular Probes) and imaged by structured illumination microscopy using a Zeiss Axio Imager M.2 with ApoTome.2 attachments and an Axiocam 506 monochrome camera. Three dimensional reconstructions were generated using Zeiss ZEN software from ApoTome image stacks ranging 10µm to 15µm depth. For labeling cryosections, this labeling protocol was modified by removing Tween-20 from blocking and washing solutions. Primary antibodies used in this study were: Rat anti-CD44 (1:1000, PFA, DSHB 5D2-27), Goat anti-FABP7 (1:1000, PFA, R&D Systems AF3166), Rat anti-Ki-67 (1:1000, PFA, Invitrogen 740008T), Rabbit anti-NFH (1:1000, PFA, Millipore AB1989), Chicken anti-NFH (1:500, PFA, Aves NFH), Goat anti-OCM (N-19) (1:250, PFA, Santa Cruz sc-7466), Rabbit anti-P75 (1:900, PFA, Millipore Sigma 07-476), Goat anti-PROX1 (1:300, PFA, R&D Systems AF2727), Rabbit anti-PROX1 (1:1000, PFA, Reliatech 102-PA32AG), Goat anti-SOX2 (1:200, PFA, Santa Cruz, 17320), Rabbit anti-SOX2 (1:1000, PFA, Millipore AB5603), Mouse anti-β1β2 TUBULIN (1:500, PFA, Sigma T8535), Rabbit anti-VANGL2 (1:1000, EMD Millipore ABN2242). Rat anti-ZO-1 (1:1000, TCA, DSHB R26.4C).

### EdU incorporation

EdU was administered to the pregnant dam by intraperitoneal injection at a dose of 10ug/g twice daily at 10am and 4pm at E16.5, E17.5 and E18.5. Tissues were collected at P0 at 4°C using 4% paraformaldehyde (PFA) in 67 mM Sorensons’ phosphate buffer (pH 7.4). Incorporated EdU was detected and labeled using the Click-iT™ EdU Cell Proliferation Kit for Imaging (Invitrogen C10337) and processed for immunofluorescent labeling to detect other proteins as described.

### Quantification

To determine cochlear length, wholemount tissue samples were collected at postnatal day 2 (P2) and stained using PROX1 antibody to label the sensory epithelium and the length of the cochlea measured using FIJI(ImageJ). Numbers of hair cells, supporting cells and EdU-labeled cells were determined using the ImageJ cell counter plugin. Counts were normalized to 100 microns of cochlear length to allow direct comparisons between different experimental groups at E17.5, P0, and P2 developmental ages.

### RNAseq analysis

Temporal bones were removed from *Emx2*-Cre; *Prox1* cKOs and littermate controls at P0 and stored at 4°C in RNAlater (ThermoFisher AM7020). Following genotyping, individual cochleae were micro-dissected to separate the organ of Corti from the spiral ganglion, outer sulcus and surrounding mesenchyme, and pooled in groups of 6 to 8 based on genotype. Tissue was frozen in liquid nitrogen and stored at -80°C until RNA extraction using Qiagen RNeasy kit (QIAGEN 74104). cDNA libraries for control and *Emx2*-Cre; *Prox1* cKOs samples (N=4 pools each) where prepared from total RNA after rRNA depletion (NEBNext Ultra II) and paired-end sequenced (150x150 bp, NovaSeq S4) at an average depth of 55.6 ± 6.6 (mean ± std) million reads per sample. Sequencing reads were inspected for quality (FastQC version 0.12.1), trimmed to 120bp to remove poly-G sequencing adapters (Cutadapt version 4.3; https://doi.org/10.14806/ej.17.1.200), and aligned (HISAT2 version 2.2.1) (Kim, Langmead et al. 2015) to mouse genome build GRCm39 (Genome Reference Consortium). On average, 34.8 ± 4.2 million reads per sample (62.6 ± 4.4%; mean ± std) successfully aligned. Differential gene expression between wildtype and cKO samples was determined using the R packages Rsubread (Liao, Smyth et al. 2019), featureCounts (Liao, Smyth et al. 2014) and DESeq2 (Love, Huber et al. 2014). To assay *Prox1* targeting, the percentage of *Prox1* reads from each sample containing the targeted exons was determined using the R package Bioconductor/Biostrings. Exons 3 and 4 sequences were split into 10 bp contigs as query, and the binary aligned sequence map (BAM) of each sample as library.

Bulk RNA-seq data from this study were compared to scRNA-seq data accessed using the gEAR interface (Kolla, Kelly et al. 2020, Orvis, Gottfried et al. 2021). Individual cell RNA profiles were identified and grouped as Hair Cells (1,035 cells) based on expression of the hair cell gene set (Ccer2, Pcp4, Cib2, Pvalb,Acbd7) or general hair cell markers (Myo6, Myo7a). Prox1 Supporting Cells (641 cells) were identified and grouped based on expression of Deiters’ cell 1/2 genes (Hes5, Pdzk1ip1, Ppp1r2, S100a1, Serpine2), Deiters’ cell 3 genes (Hes5, Igfbpl1, Lfng, Prss23, S100a1), IPC genes (Cryab, Emid1, Igfbpl1, Npy, S100b) or OPC genes (Fam159b/Shisal2b, Ppp1r2, S100b, Serpine2 and Smagp). All remaining cells (12,393) had RNA profiles that did not completely match these gene sets. For these groups, the normalized expression value of each gene per cell was summed and those values used to calculate the percentage of overall expression of each gene occurring within the group. The denominator for this calculation was the expression sum for each gene across all cells in the dataset (14,043). That value is plotted as ‘per cell expression’, with the hue of the color ranging from white (0) to saturated (1). This analysis was restricted to genes with significant changes in gene expression (padj>0.05).

